# Real-Time Spatiotemporal Tracking of Infectious Outbreaks in Confined Environments with a Host-Pathogen Agent-Based System

**DOI:** 10.1101/2024.10.01.616085

**Authors:** Suhas Srinivasan, Jeffrey King, Andres Colubri, Dmitry Korkin

## Abstract

Deadly infection outbreaks in confined spaces, whether it is a COVID-19 outbreak on a cruise ship or at a veteran home, or measles and stomach flu outbreaks in schools, can be characterized by their rapid spread due to the abundance of common spaces, shared airways, and high population density. Preventing future infectious outbreaks and developing efficient mitigation protocols can benefit from advanced computational modeling approaches. Here, we developed an agent-based modeling approach to study the spatiotemporal dynamics of an infection outbreak in a confined environment caused by a specific pathogen and to determine effective infection containment protocols. The approach integrates the 3D geographic information system of a confined environment, the behavior of the hosts, key biological parameters about the pathogen obtained from the experimental data, and the general mechanics of host-pathogen and pathogen-fomite interactions. To assess our approach, we applied it to the historical data of infectious outbreak caused by norovirus, H1N1 influenza A, and SARS-CoV-2 viruses. As a result our model was able to accurately predict the number of infections per day, correctly identify the day when the CDC vessel sanitation protocol would be triggered, single out key biological parameters affecting the infection spread, and propose important changes to existing containment protocols, specific for different pathogens. This research not only contributes to our understanding of infection spread and containment in cruise ships but also offers insights applicable to other similar confined settings, such as nursing homes, schools, and hospitals. By providing a robust framework for real-time outbreak modeling, this study proposes new, more effective containment protocols and enhances our preparedness for managing infectious diseases and emerging pathogens in confined environments.

## Introduction

Infection outbreaks affecting people in predominantly confined environments, such as schools, nursing homes, college dormitories, large open-space offices, meat packing facilities, grocery stores, and cruise ships, present a substantial healthcare challenge due to increase in infection risks and often limited access to comprehensive medical care facilities and ask for new construction policies and guidelines (*1-5*). The COVID-19 pandemic further emphasized the critical need to understand the dynamics of infectious outbreaks due to devastating outbreaks on Diamond Princess cruise ship (*6*), USS Air Career Roosevelt (*7*), and at Kirkland, WA and Holyoke, MA nursing homes (*8, 9*). Cruise ships have been at the epicenter of main infectious outbreaks, caused primarily by noroviruses (*10*), influenza (*11*), and lately SARS-COV2 (*12*), due to increased frequency of close interactions between a large cohort of individuals, and the presence of high foot traffic areas, such as dining rooms and theaters, thus being one of the best documented sources of historic data on infection dynamics (*13*). Unfortunately, the current containment protocols developed primarily for norovirus infections proved not to be efficient for SARS-CoV-2, perhaps due to the extended asymptomatic period with viral shedding and large percentage of asymptomatic carriers infected with COVID-19 (*14*) and its different primary mode of transmission (airborne for SARS-CoV-2 and surface-based for norovirus) (*15, 16*).

The ability to track the spatiotemporal dynamics of an infectious outbreak in a confined environment in real-time is critical for determining key factors driving the outbreak and developing optimal protocols for its effective and efficient containment. Conventional epidemiological models (*17-19*), including the ones recently developed to study transmission dynamics of COVID-19 (*20*), are large-scale, providing an averaged population-level description and unable to integrate complex details on the individual interactions between the environment, host and pathogen, such as person-to-person transmission, population density variation within a contained space, pathogen transmission characteristics. In addition to the conventional compartmental epidemiological models (*21*), an artificial intelligence (AI) approach, agent-based models (ABMs), have been recently applied in epidemiology and public health settings (*22, 23*). Notably, ABMs can represent the interaction and heterogeneous behavior of individuals (agents) within a system rather than the aggregate behavior of populations in compartmental models. Aggregate behavior then becomes an emergent property of ABM systems, providing increased accuracy as compared to other modeling systems (*24, 25*). The focus on individual behavior of the agents also allows for the ability to scale ABMs, from focusing on individual transmission dynamics (*21*), to urban contexts (*26, 27*), and even country-wide epidemic modeling (*28-30*). However, such models have not incorporated detailed host behavior in a specific environment (*31*) and importantly, no epidemic model to date has used a highly detailed three-dimensional environment representation which is traversable by both, the host agents and pathogens (*32*). Here, we develop a new approach to study real-time spatiotemporal dynamics of an infection on cruise ships by utilizing an ABM framework, pathogen modeling, and geographic information system (GIS) representation of the cruise ships. We first assess this approach using historic data from past outbreaks of norovirus, whose economic and health burden is well-documented (Supp. Fig. S1). Then, we analyze the dynamics of potential outbreaks for SARS-CoV-2, Ebola virus, and 2009 H1N1 pandemic influenza virus, identifying the pathogen-specific key contributing factors and suggesting improvements to the current containment protocols.

## Results

In our agent-based modeling approach, called AI-GIS Infection Dynamics (AGID), a real-world cruise ship (which we refer to as Ship X) was represented as a three-dimensional geographic information system (GIS) through modeling each deck as a two-dimensional GIS and connecting the decks into a spatially ordered network, where connections corresponded to the elevators and stairs between the decks (Fig. 1 and Supp. Methods, Section 1). The hosts, *i*.*e*., passengers and crew members, were modeled as agents which allowed to explicitly define complex interactions between the hosts, between hosts and pathogens, as well as between hosts and environment. The two classes of agents, passengers, and crew, each have specific parameters defining their behavior in the system, which were then combined with pathogen modeling (Fig. 1A). Pathogens were modeled by defining the probability of infection and illness for each exposed agent based on dose-response in real-time.

**Figure 1.**
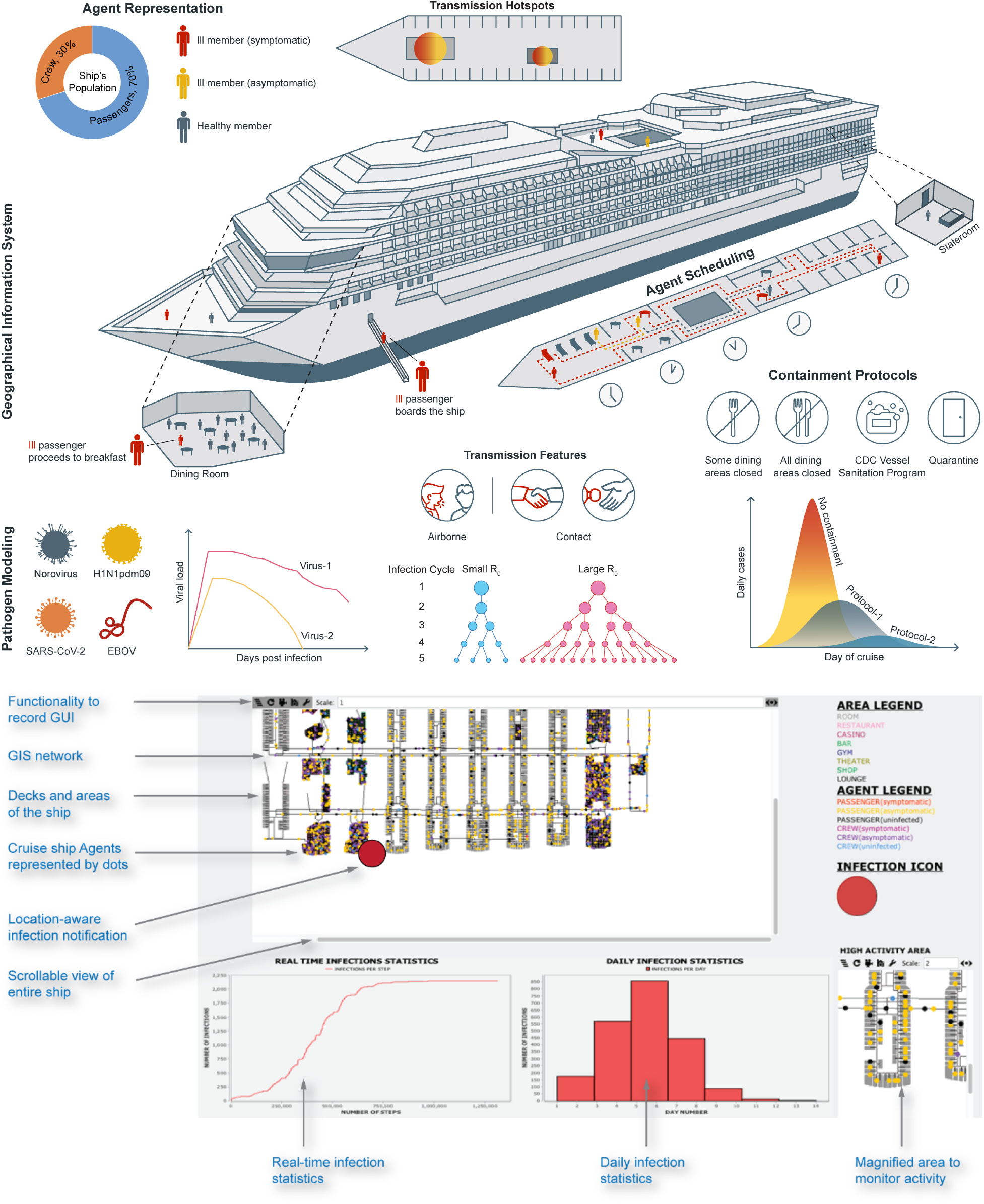
Overview of the AGID real-time infection modeling system. **(a)** The core of the system consists of the geographical information system (GIS) based 3D representation of the cruise ship as a traversable network for the agents. Agent modeling consists of a population representation and action modeling where the crew and passenger agents follow their respective schedules. Pathogen modeling is carried out individually for each viral strain through a set of parameters, such as viral shedding, incubation period and R_0_ among others. CDC vessel sanitation program and other containment protocols are implemented to study their effects in mitigating an outbreak. (**b**) The graphical user interface (UI) provides real-time display of agent movement, pathogen transmission and infection cases. The UI provides functionality to interact with the GIS by zooming in/out into high-density areas. The cruise ship areas and agents states are represented by a colored legend. The infection icon is displayed specific to the location where a previously infected agent successfully passed the infection to a healthy agent.

### Assessment of critical parameters affecting infection dynamics

We first assessed the importance of explicitly defining in an AGID model the agent-specific parameters. We compared the performances of three basic—naïve—types (Types I, II & III, Fig. 2A, Supp. Fig. S2) of models, where the key host and/or pathogen parameters were substituted by altered values, not reflecting the real-world characteristics. Eight naïve models, which we referred to as Naïve Sim-1 – Naïve Sim-8, were developed to simulate a specific norovirus outbreak that occurred on Cruise Ship X in 2006, and assessed by comparing the simulations results against the historical data (Fig. 2A and Supp. Fig. S2). The historically observed cases over the nine-day cruise reached an attack rate (AR) of 6.5%. The models of Type-I (Naïve Sim-1 & 2) were designed to assess the importance of explicitly specifying the daily routines for the host agents. Naïve Sim-1 lacked the information on the daily activities of the host agents: instead, the agents randomly traversed the ship without any breaks, while other parameters, including pathogen-specific characteristics, were accurately defined (Supp. Fig. S2). Since an infectious agent did not spend significant time in any location or interacted with another host agent, the infection spread was substantially lower compared to the historical data, with AR of ∼1%. Adding the breakfast schedule in Naïve Sim-2 increased agent interaction in dining rooms, causing AR to reach 2.5%. In models of Type-II (Naïve Sim-3, 4 & 5) the agents still performed a randomized traversal of the ship, while being infectious with the higher, and therefore more impactful for the infection, characteristics. Specifically, in Naïve Sim-3 the agents were infectious all the time; in Naïve Sim-4 an infectious agent transmitted the pathogen to all agents in the vicinity; and in Naïve Sim-5 the agents were infectious all the time and infect everyone in the vicinity.

**Figure 2.**
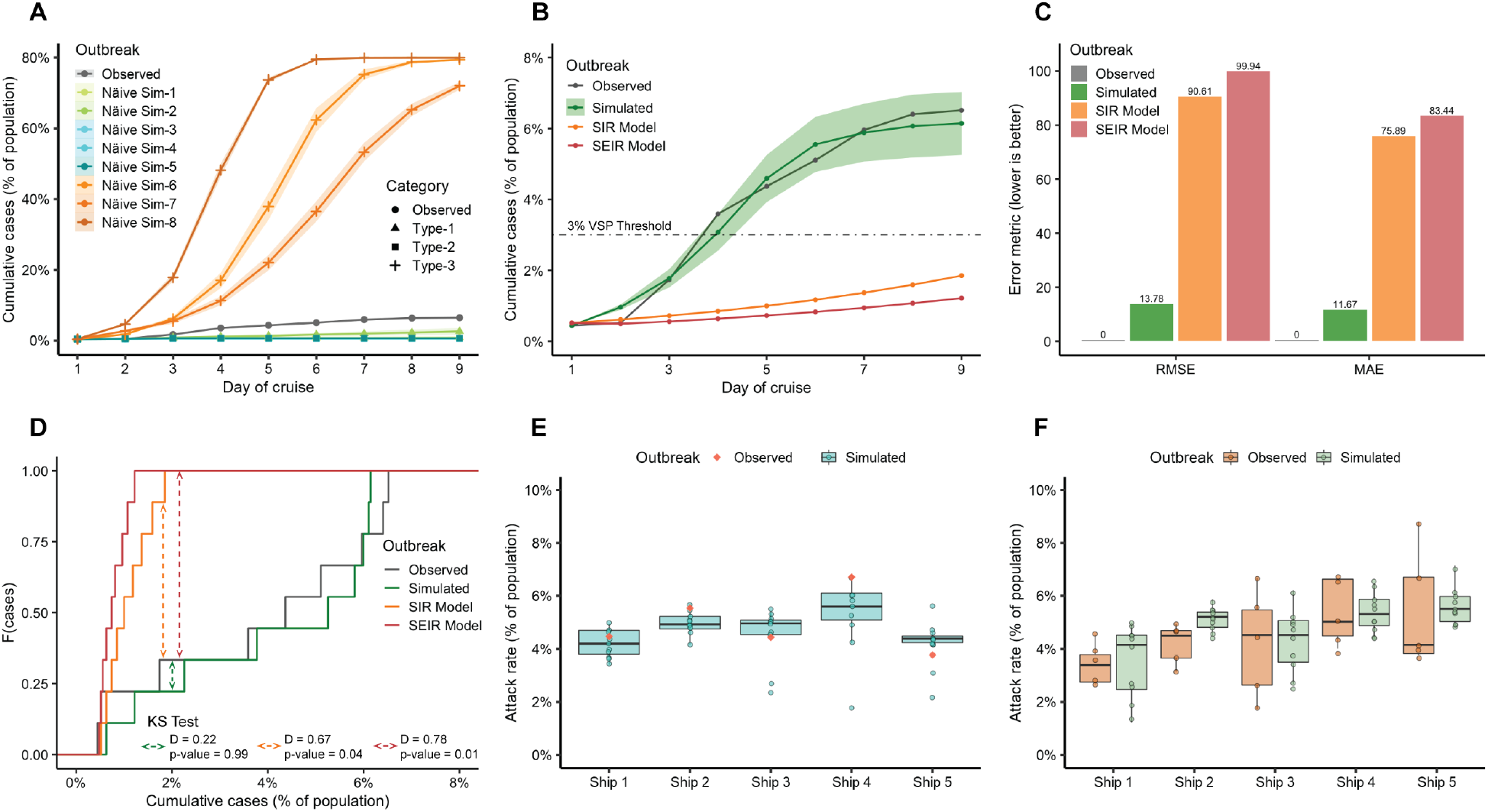
Evaluation of simulated norovirus outbreaks shows a superior performance by AGID models. (**A**) Simulation of three types of naïve agent models and pathogen modeling (total of eight different set-ups), and comparison with the observed outbreak on Ship X. The naïve models failed to match the observed outbreak. (**B**) The detailed agent and pathogen modeling in AGID (green, depicted with the standard deviation) is closely aligned with the observed epidemic curve on Ship X (grey) w.r.t cases and time, including reaching VSP threshold, whereas the conventional SIR (orange) and SEIR (red) model predictions differed substantially from the observations. (**C**) Comparative assessment of the prediction by the AGID model simulations (green) together with conventional SIR (orange) and SEIR (red) models w.r.t historical observations (grey); both RMSE and MAE were an order of magnitude greater for the two conventional models. (**D**) Kolmogorov–Smirnov test of the predicted and historically observed cases showed the predictions from AGID simulations (green) were similar to the distribution of the observed cases (grey), with a small D value, while SIR (orange) and SEIR (red) predictions had large D values and did not satisfy the null hypothesis. (**E**) Ten AGID simulations for each of the five ships were performed to validate the system generalizability by recreating specific outbreaks observed on the five cruise ships showed that most predictions had less than one percent deviation from observed outbreak. (**F**) AGID simulations (light green) to validate the system’s variability in predictions by providing only median parameters for the five ships, and comparing against the variation in observed outbreaks (light orange): the mean values are similar and mostly withing a percentage point.

Increasing the transmission rate did not increase AR beyond 1% (Supp. Fig. S2), because the agent-to-agent interaction was still minimal. Models of Type-III explored the impact of the parameters defining the infection transmission alone. Specifically, in three models, Naïve Sim-6, 7 & 8, the host agents followed an accurate schedule, while the parameters accounting for pathogen transmission were changed in the same way as in Naïve Sim-3, 4 & 5, respectively. In a stark contrast to the previous five models, here for Naïve Sim-6, 7, 8, we observed a substantially higher AR, compared to the historical data, reaching a maximum of 80%. The infection numbers increased sharply due to the higher than usual transmission dynamics from Naïve Sims-6–8 (Fig. 2A). Overall, none of the eight naïve simulations matched the observed outbreak: they were substantially different at each observed time point, demonstrating the critical needs of the parameters describing both, host actions and pathogen transmission, to accurately model an outbreak.

### Evaluation of AGID model and comparison with the conventional models

Next, once all parameters of the AGID model were defined, we compared its simulation accuracy to the accuracies of two conventional compartmental models of infection outbreaks, SIR (susceptible, infectious or recovered) (*33*) and SEIR (susceptible, exposed, infectious or recovered) (*34*), recently used for modeling COVID-19 outbreaks in nursing homes (*35*). All models were compared to the real-world data, revealing that AGID model’s predictions closely matched the trends in the observed outbreak (Fig. 2B) with respect to the number of cases per day. The AGID simulations also reached the 3% CDC VSP threshold on Day 4, consistent with the observed data, which indicated that 3% threshold was met after ∼3.5 days. The SIR and SEIR predictions did not reach the 3% threshold over the entire cruise duration. Furthermore, both observed and AGID-simulated cases plateaued at Days 8 and 9, reaching a total attack rate of 6.25% and 6.5% respectively. When comparing the performance of the AGID simulations against SIR and SAIR using RMSE and MAE measures (Fig. 2C), we observed that the AGID simulations achieved the lowest RMSE (13.78) and MAE (11.67) values, an order of magnitude lower than the corresponding values for the SIR (90.61 and 75.89) and SEIR (99.94 and 83.44) models. As the real and predicted outbreaks were each representative of the case probability distribution over the nine-day cruise period, we used the Kolmogorov-Smirnov (K-S) test to quantify the difference in the predictions. The three probability distributions were plotted as a function of the cumulative cases (Fig. 2D), and we found that the AGID simulations were similar to the observed outbreak, while the SIR and SEIR models provided poor prediction forecast. The K-S test statistic on the distance between the distributions was the smallest for the AGID simulations (0.22), but substantially larger for the SIR and SEIR models, 0.67 and 0.78 respectively, with the statistical significance (*p* < 0.05).

The evaluation was then expanded to test the ability of the AGID simulations to accurately predict multiple outbreaks across five different cruise ships. The cruise ships differed with respect to the overall layout, which includes the number of floors, staterooms, facilities and their types, as well as occupants (Supp. Table S1). The cruise ships were selected from CDC data such that (1) each ship was involved in at least five different norovirus outbreaks and (2) their layout can still be modeled using the GIS of Ship X. First, we assessed the accuracy of predicting an outbreak on each of the five ships (Fig. 2E). The respective population sizes and cruise durations were used as initialization parameters from the real-world outbreak reports to create the simulations. The results revealed the accuracy of the AGID model in predicting the attack rates across all five cruise ships. The deviation was less than 1% in the median value for the attack rates predicted by our model versus the observed attack rate (Fig. 2E). Additionally, for Ships 1–4, the observed attack rate was within the interquartile range of the simulated attack rate. Next, we further expanded our evaluation to model five independent outbreaks for each of the five cruise ships: we expected to see that each ship had a defined range of attack rates if the GIS, population, and pathogen are maintained as constant variables. For this purpose, we use the mean values for each ship to seed the simulations. We analyzed the distributions of observed and simulated attack rates (Fig. 2F) and found that the median values were highly similar with a difference of less than 1% for Ships 1–4 and ∼1.5% for Ship 5.

### Application of AGID to simulate infectious outbreaks across different pathogens

We next applied the AGID model to study other types of infectious outbreaks on cruise ships which had previously occurred or considered a possibility (Fig. 3, Supp. Figs. S3, S4). We focused on the infections caused by viruses that recently emerged: norovirus, H1N1 2009 pandemic influenza A strain (H1N1pdm09) (*36*), SARS-CoV-2 virus causing the recent COVID-19 pandemic (*37, 38*), as well as Zaire ebolavirus (EBOV), which was linked to more than two dozen outbreaks in Africa and a recent 2014 spread in U.S. and Europe (*39*). The viral pathogens were selected such that we could analyze the variation in transmission modes, disease cycles, attack rates, fatality rates, and the impact of novel pathogens. The shedding values for H1N1pdm09 and EBOV were obtained from animal model studies (*40, 41*), the values for norovirus were obtained from the human challenge studies (*42*), while the values for SARS-CoV-2 (original strain) were obtained from clinical testing (*43*) (Supp. Table S2 and Supp. Fig. S5). Two different transmission modes were introduced into our model: airborne (direct transmission) and contact or surface-based. The airborne viruses, H1N1pdm09 and SARS-CoV-2, have a significant proportion of asymptomatic cases, 10% and 50%, respectively (*14*, 15, 44, 45), making the containment more challenging (Fig. 3A). On the other hand, norovirus and EBOV have a very low or unlikely chances of asymptomatic carriers (*46*). In addition, to the original strain of SARS-CoV-2, we also simulated the infection dynamics of the Delta variant, which was one of the Variants of Concern being monitored by the CDC, due to its higher transmissibility and decreased neutralization by antibodies (*47*).

**Figure 3.**
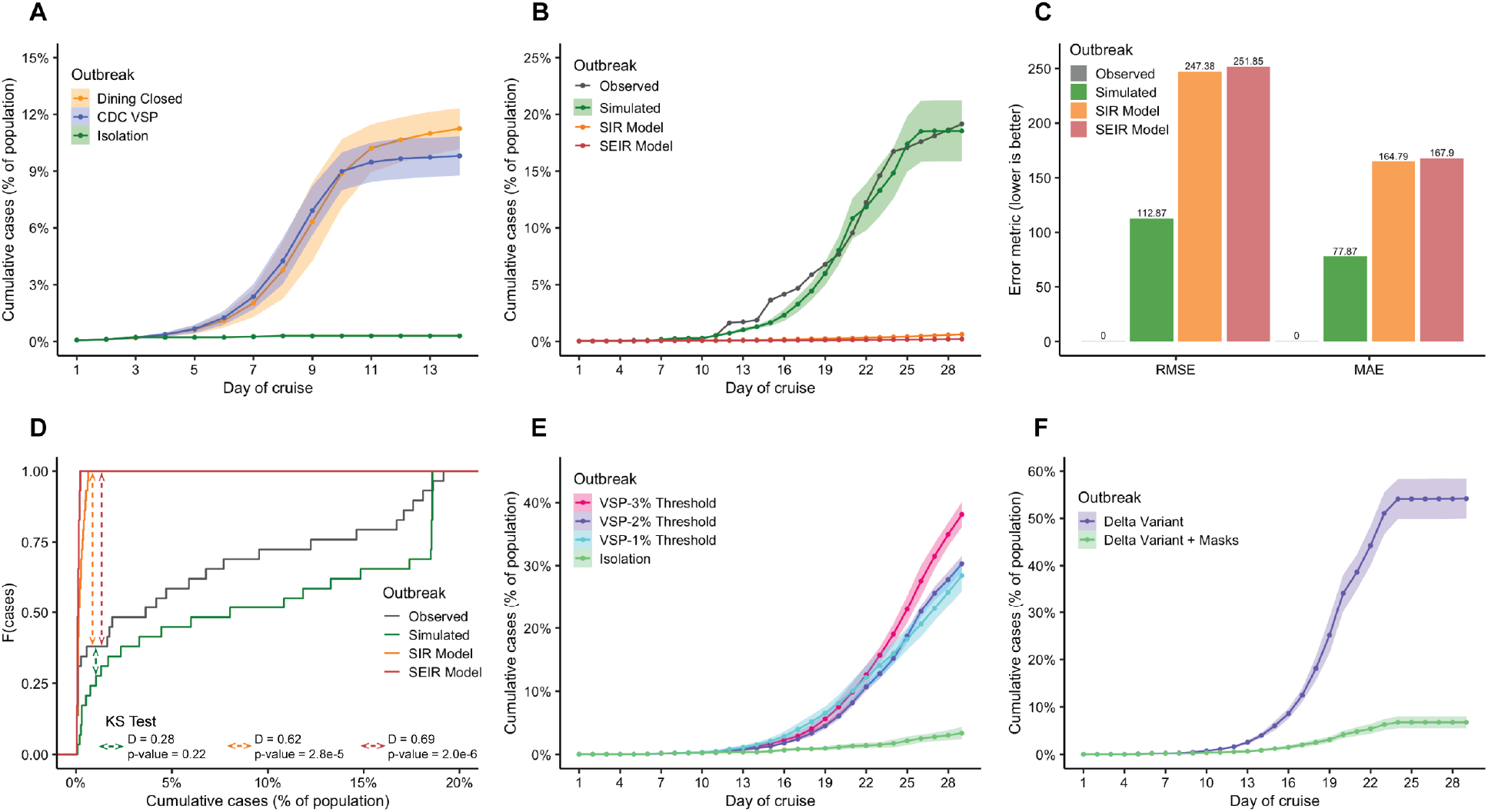
Simulation of H1N1 and SARS-CoV-2 outbreaks, including Delta variant and efficiency of the containment protocols. (**A**) AGID simulation of H1N1pdm09 virus strain highlights the importance of implementing pathogen-specific protocols, with Isolation (green) being the only effective protocol for that outbreak, while closing the dining areas (orange) and implementing CDC VSP (blue) did not substantially reduce the number of infections. (**B**) AGID simulation (green, depicted with the standard deviation) of the original strain of SARS-CoV-2 using Ship X GIS representation, with conventional SIR (orange) and SEIR (red) models and comparison with the historically observed cases (grey) on the Diamond Princess cruise ship. (**C**) Errors present in the prediction of the AGID simulations (green), SIR (orange) and SEIR (red) models w.r.t Diamond Princess observations; both RMSE and MAE are greater by a factor of two for the SIR and SEIR models. (**D**) Kolmogorov–Smirnov (KS) test of the predicted cases and observed cases, showed the AGID’s predictions (green) are more similar to observed case distribution (grey) than the prediction by SIR (orange) and SEIR (red) models, with a smaller D value. (**E**) Simulations show that lowering VSP threshold from 3% (red) to 2% (purple) and even to 1% (blue) for SARS-CoV-2 (original strain) do not provide better containment, while the Isolation protocol (green) is more effective. (**F**) AGID simulations show the benefit offered by face masks (green) to reduce the case load, as compared to the “no-mask” infection dynamics (purple).

The AGID simulations for SARS-CoV-2 transmission dynamics accurately modeled the infection trajectory relative to the population and number of days (18.6% attack rate), with a similar distribution of the case load to that observed on the Diamond Princess cruise ship (Fig. 3B). Our model estimated that the Diamond Princess COVID-19 outbreak had an average incubation period of approximately five days, consistent with the previous reports (*48*). Compared to the compartmental models, the simulations were more accurate as demonstrated by the error metrics, RMSE of 112.87 and MAE of 77.87 for simulations, and the SIR and SEIR predictions have both RMSE and MAE greater than double the error metrics of the simulations (Fig. 3C). Additionally, the K-S test statistic on the distance between the distributions is smallest for the simulations (0.22), but substantially larger for the SIR and SEIR models, 0.62 and 0.69 respectively (Fig. 3D), with the statistical significance (*p* < 0.05). The model highlighted the importance of the significant proportion of asymptomatic carriers in driving the outbreak. Interestingly, when we applied AGID model to simulate the infection dynamics of the Delta variant, the simulation achieved an attack rate of 54.2%, nearly three times as high as the original virus strain (Fig. 3F).

The spatiotemporal analysis of the infection dynamics using the simulation data from the AGID models demonstrated that the high-density and high foot traffic areas were the infection hubs, where the pathogen transmission occurring most frequently for any virus (Figure 4, Supp. Figs S7, S8). The comparison of different viral infections in the same confined environments suggested that the mode of transmission of the virus played a key role in determining the rate of infection. In particular, the comparative analysis of infection dynamics for norovirus and the original strain of SARS-CoV-2 revealed an important difference between the transmission hotspots during the outbreaks (Figure 4). While both outbreaks had a common hotspot—a hallway leading to a theater (Supp. Fig. S6), where the regularly scheduled events were held— the norovirus infection-specific hotspots were the dining areas, where the hosts would be seated at a distance from each other, but would nevertheless interact with the fomites (*e*.*g*., tables), possibly contaminating it and ensuring host-fomite-host mode of transmission, which is characteristic for norovirus infections (*49*). In addition to the transmission mode, by comparing the four viruses, Norovirus, 2009 H1N1 pandemic influenza, SARS-CoV-2, and Ebola, we observed that the incubation period was also a major determinant of the attack rate achieved during the duration of a cruise. EBOV had a long, 8–10 day incubation period, resulting in low number of infection cases during the cruise (Supp. Fig. S4), whereas for norovirus it was a substantially shorter incubation period, 1–3 days, which resulted in more cases (Fig. 2B). When studying the COVID-19 outbreaks, characterized by an incubation period of approximately 5–7 days, we observed that the initial growth rate for the original strain of SARS-CoV-2 was slower compared to the norovirus. However, it eventually followed an exponential trend, driven by its mode of transmission. Lastly, the impact of asymptomatic carriers on the spread of infection varied significantly across different viral pathogens: norovirus exhibited negligible asymptomatic transmission, H1N1pdm09 had approximately 10% of cases transmitted asymptomatically, while the original strain of SARS-CoV-2 involved nearly 50% asymptomatic carriers.

**Figure 4.**
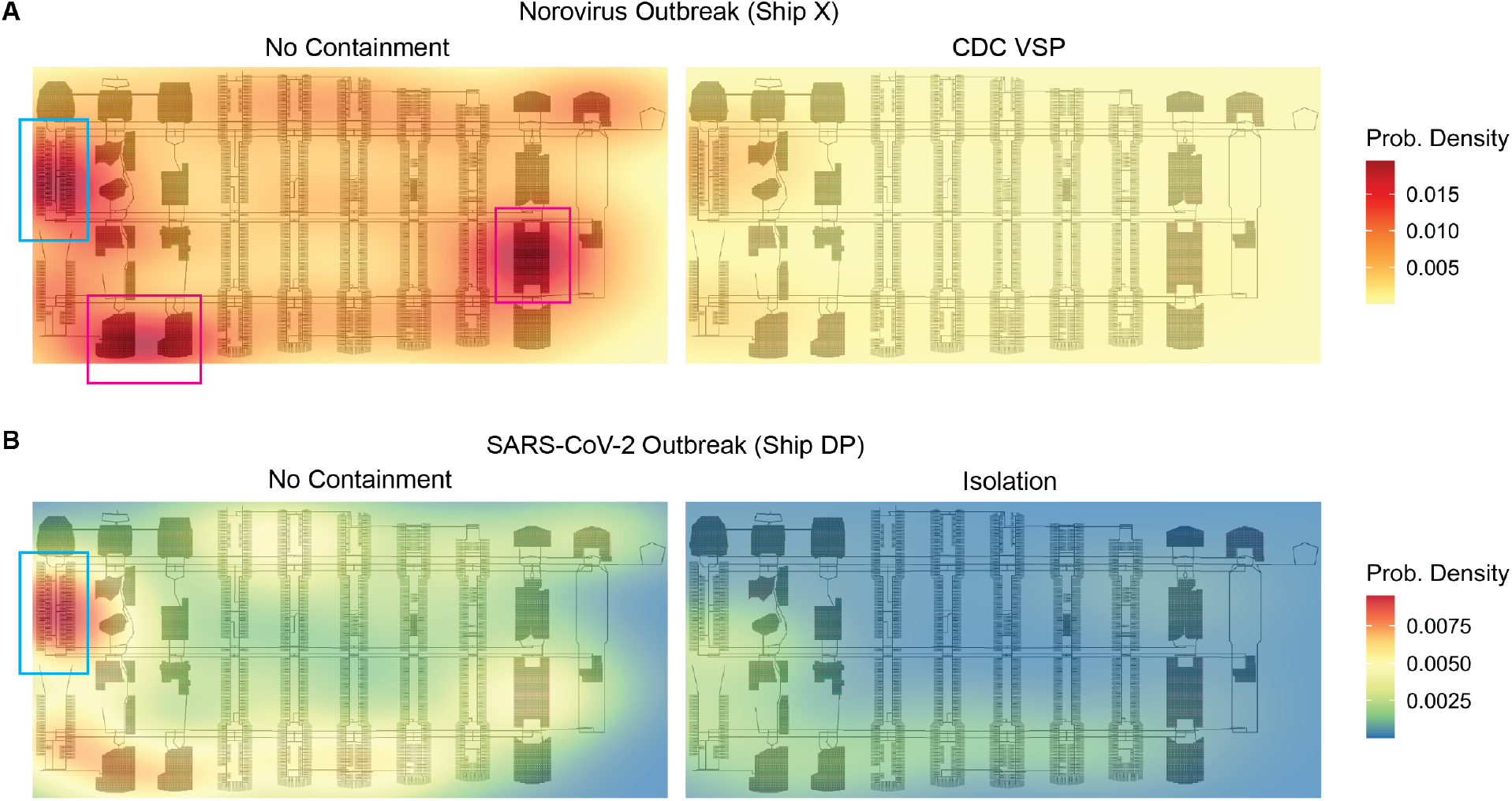
Spatial distribution of infection hotspots and efficacy of containment protocols. (**A**) Simulation of a Norovirus outbreaks on Ship X during a nine-day cruise. The left panel showing the infection hotspots for a No Containment scenario (AR: 39%) with primary concentration in the dining halls and the right panel showing the effectiveness of CDC VSP to reduce transmission (AR: 6%) (color scale shows two-dimensional kernel density estimate for hotspots). (**B**) Simulation of SARS-CoV-2 (wild type) outbreaks on the ship Diamond Princess (Ship DP) during a 30-day cruise, with the left panel showing the outbreak based on the documented 2020 outbreak (AR: 19%) and the right panel showing implementation of Isolation procedures that largely reduces transmission (AR: 4%) (color scale shows two-dimensional kernel density estimate for hotspots). The outbreaks for both viral pathogens create well-defined infection hotspots when no containment protocols are implemented. We note that the distribution of the infection hotspots are different and dependent on the transmission mode: for norovirus the most prominent hotspots are the restaurants (red squares) and the main crew corridor (blue square), whereas for SARS-CoV-2, it is only the main crew corridor. The restaurants are modeled to have regularly scheduled events for a large number of agents, while the main crew corridor has the highest traffic among the crew members.

### Containment protocols: Rethinking existing measures through an AI-driven optimization

One of the main utilities of the developed AGID approach lies in its potential to optimize the existing and propose new pathogen-specific containment protocols. The obtained AGID model has showed effectivity of the CDC-defined outbreak containment measures for norovirus (Fig. 4A), which are currently triggered when attack rate reaches 3% (*50*). It also showed that another containment protocol, *i*.*e*., closing all dining areas (Dining Closed) was nearly as effective in controlling the norovirus outbreak: the attack rates for CDC VSP and for the Dining Closed protocol were ∼6.5% and ∼7%, respectively (Supp. Fig. S3). The GIS framework enabled the spatial analysis of hotspots and helped understand a striking phenomenon—a rapid increase in the number of infection cases during the gradual closure of the dining areas. For instance, when only a single dining area was closed the outbreak size (Supp. Fig. S3, Dining Restricted) surpassed that one for no containment scenario (Supp. Fig. S3, No Containment). Indeed the partial restriction of the dining areas worsens the overcrowding effect in the remaining dining rooms (Supp. Fig. S9), thus leading to the increased infection rate.

Furthermore, for the SARS-CoV-2 virus (original strain), lowering the VSP threshold from 3% to 2%, and even to 1% was not sufficient to mitigate the outbreak (AR were 53.7%, 45.7%, and 42% respectively). Additional restrictions on activities or the use of the face masks were determined to be necessary to reduce the case load. While the lowered thresholds for VSP were not efficient, we found that the passenger isolation on presentation of symptoms for the original strain (AR: 5.7%) and face masks on symptom presentation for the delta variant (AR: 6.7%) were able to substantially reduce the case load (Fig. 3E and 3F). We also found that the Isolation protocol was efficient when simulating an Ebola outbreak, due to the long and varying incubation periods, differing symptoms, and initial small case load, all of which would not meet the criteria for VSP (Supp. Fig. S4). Overall, the spatiotemporal analysis identified common shifts in the patterns of pathogen transmission: from the explosive nature of infection in the high-traffic areas, such as main crew corridor, when no containment protocol was implemented, to the localized infections in the staterooms with the Isolation protocol implemented, where only individuals in close physical proximity were infected (Fig. 4B).

### Availability

The simulation source code is freely available at https://github.com/KorkinLab/infection-dynamics

## Materials and methods

### AI-GIS Infection Dynamics (AGID) model

The AGID-based model for a specific pathogen in a confined space is dependent on the space’s environment. For a cruise ship, the environment is characterized by the number of decks and their dimensions, number and spatial distribution of staterooms, dining rooms, other recreational facilities, hallways, stairs, and elevators (Fig. 1B, Supp. Fig S6, Supp. Methods, Section 1.1). The AGID model is also dependent on the number of index cases. The agents (hosts) include passengers and crew members; these two groups are primarily distinguished by their daily schedules (Supp. Figs. S10, S11, and S12, Supp. Methods, Section 1.2). A host agent is also characterized by several time-dependent parameters, such as the current health state, daily activity status, and viral load. In AGID model, we distinguish five health states for a host agent: healthy, infected asymptomatic, infected symptomatic, recovered, and perished. The transition between the states (Supp. Fig. S13) is determined based on the average timeline of the disease life cycle extracted from literature (Supp. Table S2). A pathogen model is defined for a specific infection and is characterized by the basic reproduction number (R_0_), quantified transmission modes, incubation period, viral shedding curve, and fatality rate (see Supp. Methods, Section 1.3 for more detail). The viral exposure by a susceptible host is determined by the number of viral particles from another infected host that is in proximity to the susceptible host and calculating the probability of being infected when exposed to the virus:

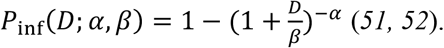

The probability of the infected host to progress from incubation period to illness stage is then estimated as *h(D*|*η, r) =* 1 − (1 + *ηD*)^− *r*^, where *D* is the dose of viral particles, *α* and *β* are the infectivity parameters, and *η* and *r* are the maximum-likelihood estimates obtained from literature (*53*).

The simulation of a viral outbreak is performed by first setting the initial number of infected passengers and the maximum allowed onboard population and then running the simulation for a specified duration defined by the length of the cruise. During the simulation, the host agents perform a sequence of daily routines, with starting times of recreational activities having variation based on occupancy of common areas on the ship (Supp. Figs. S10 and S11). For the infected and symptomatic crew members, we also implement a commonly used crew containment protocol (Supp. Fig. S12). In the simulation, we explicitly implement an interaction between a pair of host agents, one of which being infected, by defining the interaction between two agents based on their proximity and duration, specifying the number of particles transmitted through air or surface, and determining the probabilities of infection and then illness for the host agent exposed to the pathogen. We used the simulation framework MASON to implement our agent-based model and perform the simulations (*54*), and the GIS was constructed using ArcGIS (*55*), the graphical user interface (GUI) (Fig. 1B, Supp. Methods, Section 1.1) was developed using Java Swing (*56*).

### Evaluation protocol

The accuracy of our model is assessed using historic cruise ship outbreak data available from the Centers of Disease Control and Prevention (CDC), as well as by comparing the model against several naïve, random simulation models and against conventional compartmental models of infections (Supp. Methods, Section 2). First, we evaluate if the proposed system can accurately simulate a norovirus outbreak on a particular cruise ship. For the cruise ship X, we collect detailed daily infection data for an outbreak that happened over a nine-day period in 2006. Second, we evaluate the generalizability of the system for the same pathogen (norovirus) in five additional cruise ships of various physical sizes and population sizes by following a rigorous protocol consisting of two steps: (i) modeling a specific recorded norovirus outbreak on each ship (ii) modeling a typical or averaged norovirus spread by aggregating five recorded outbreaks on each ship (Supp. Methods, Section 2 and Supp. Table S1). The CDC Vessel Sanitation Program (VSP) (*50*) is implemented for each ship using standard guidelines, *i*.*e*., when the number of infected individuals reaches 3% of the cruise ship population. The CDC reports provide the total number of passengers and crew, and the total number of infections reported for the duration of the outbreak. We also assessed the accuracy of the approach’s agent and pathogen modeling by creating three types of indiscriminate/naïve models, and using SIR (susceptible, infectious or recovered) and SEIR (susceptible, exposed, infectious or recovered) models to predict the outbreak on Ship X. To systematically compare predictions of the models with the real outbreaks, we use three metrics (Supp. Methods, Section 2.2): root-mean-square error (RMSE) (*57*), mean absolute error (MAE) (*57*), and Kolmogorov-Smirnov (K-S) test (*58*). Furthermore, because of its stochastic nature, our model was evaluated for its consistency to produce robust predictions across multiple runs with the same model parameters. To ensure the computational feasibility of the above assessments, we performed 10 simulations for each combination of GIS, agent, and pathogen parameters, resulting in a total of 190 simulations.

### Description of containment protocols

We next implemented and simulated four containment strategies to determine their efficiency in limiting the spread of an infection (Supp. Table S3, Supp. Methods, Section 1.4). Containment strategies (protocols) are necessary to modify the behavior of individuals in a population to reduce the spread of an infectious outbreak to ultimately “flatten the curve” (*59*). The protocols are designed based on the environment where an outbreak occurs and are often formulated by a governing agency, such as CDC. The implemented containment protocols included: (1) closure for cleaning of a dining area where the infection was initially detected; (2) closure of all dining areas, with individual meals delivered to staterooms; (3) CDC vessel sanitation program (VSP), a protocol developed for infections with gastrointestinal illness symptoms, initiated when 3% of passengers or crew report the symptoms; and (4) isolation, where passengers voluntarily isolate themselves when symptoms appear and continue the isolation until symptom free. Additionally, for the evolving dynamics of COVID-19, we implemented two other containment protocols, including the lower VSP thresholds and mandatory wearing of the face masks (Supp. Methods, Section 1.4).

## Conclusions

In summary, our study demonstrates that an agent-based modeling approach that integrates both host and pathogen functionalities, along with the detailed GIS-based representation of the environment, can provide unique spatiotemporal insights into the infection dynamics, as well as determine important dos and don’ts during an outbreak. The real-time AGID models developed in this work offer live visualization of the outbreak and allow studying the infection dynamics at the individual host resolution, tracking the health status and changes of each individual. These models accurately predict the impact of epidemics, measured as a function of the daily number of infected agents, for different viruses with specific virulence and infectivity rates. Notably, the AGID models predicted the attack rates with lower error at each time point, outperforming the conventional epidemiological models. The high-resolution GIS presentation reveals infection hotspots on the cruise ship, which traditional ODE-based models cannot capture. Additionally, the GIS representation facilitates the development and simulation of new containment protocols, such as closing all dining areas, to better control the viral outbreak. Our models allow for better understanding of the pathogenicity of novel viruses, including likely incubation periods, the rate and impact of asymptomatic carriers, and the effectiveness of current containment protocols against the emerging pathogens and their variants.

One of the most significant advantages of the AGID models is in their capability to systematically evaluate and compare the effectiveness of various containment protocols. Our model demonstrated that in the case of norovirus outbreaks on cruise ships, closing all dining areas, while leaving other spaces unrestricted, could significantly reduce transmission. At the same time, the model revealed that closing only one or a few main dining areas, would surprisingly have the opposite effect, creating tighter population bottlenecks in the remaining open dining rooms, which in turn increased infection rates.

Furthermore, we found that airborne influenza-like illnesses require a substantially lower threshold for initiating practically any type of containment measures, ranging from closing dining and recreational areas to isolating all passengers. This finding underscores the need for pathogen-specific containment strategies and the importance of rapid, onboard pathogen identification as part of outbreak prevention.

Our AGID model showed that containment protocols restricting the movement of symptomatic individuals were highly effective, particularly when individuals self-isolated at the first signs of symptoms or when ship-wide quarantine measures were implemented, as observed during the real-world events on Diamond Princess.

The agent-based modeling, while a powerful approach, is not without its limitations. Data parameters, such as the reproductive rate for infectious diseases, are often difficult to obtain from the literature. In addition, assessing the model’s validity can be challenging, particularly when modeling unobserved associations. Further improvements of the model’s predictive power can be done by making agents more adaptive and interacting more intelligently by leveraging recent machine learning techniques, such as reinforcement learning (*60*), and by incorporating more complex models to describe the environment (*61*), like computational fluid dynamics to accurately model air conditioning systems (*62*).

As the world recovers from the COVID-19 pandemic, it is not a question of if, but when the next global pandemic will occur (*63*). The confined environments we encounter in our everyday lives pose heightened risks for infection spread due to the close proximity of healthy and infected individuals. Computational approaches are among the most effective tools not only to forecasting the impact of such outbreaks in various social settings but also to proposing efficient strategies to mitigate them. While AGID approach was initially applied to study infection outbreaks on the cruise ships as one of the most well-documented types of confined environment, it can be adapted to virtually any other setting, including schools, grocery stores, nursing homes, or movie theaters. In the future, the model can be further enhanced to incorporate additional factors, such as vaccination status and innate immunity within the population, the implementation of hygiene measures like hand washing and mask-wearing, and the consideration of infection-specific age and gender groups.

## Acknowledgements

We thank Andi Dhroso, Andrew Lund, and Timothy Matisziw for the useful discussion and contribution to the initial version of the agent-based framework and Isabelle Cordova for designing the cruise ship infographic portion of Figure 1. We also thank Jacob Collins, Pardis Sabetti, and Stephen Shaffner for critical review of the manuscript and useful feedback.

## References

1. C. Beggs, C. Noakes, P. Sleigh, L. Fletcher, K. Siddiqi, The transmission of tuberculosis in confined spaces: an analytical review of alternative epidemiological models. The international journal of tuberculosis and lung disease 7, 1015–1026 (2003).

2. J. K. Gupta, C. H. Lin, Q. Chen, Risk assessment of airborne infectious diseases in aircraft cabins. Indoor air 22, 388–395 (2012).

3. H. Ito, S. Hanaoka, T. Kawasaki, The cruise industry and the COVID-19 outbreak. Transportation research interdisciplinary perspectives 5, 100136 (2020).

4. T. L. Gustafson et al., Measles outbreak in a fully immunized secondary-school population. New England journal of medicine 316, 771–774 (1987).

5. L. Morawska et al., A paradigm shift to combat indoor respiratory infection. Science 372, 689–691 (2021).

6. S. Tabata et al., Clinical characteristics of COVID-19 in 104 people with SARS-CoV-2 infection on the Diamond Princess cruise ship: a retrospective analysis. The Lancet Infectious Diseases 20, 1043–1050 (2020).

7. D. C. Payne et al., SARS-CoV-2 infections and serologic responses from a sample of US Navy service members—USS Theodore Roosevelt, April 2020. Morbidity and Mortality Weekly Report 69, 714 (2020).

8. P. M. Davidson, S. L. Szanton, Nursing homes and COVID-19: We can and should do better. Journal of clinical nursing, (2020).

9. S. Bagchi et al., Rates of COVID-19 among residents and staff members in nursing homes—United States, May 25–November 22, 2020. Morbidity and Mortality Weekly Report 70, 52 (2021).

10. E. T. Isakbaeva et al., Norovirus transmission on cruise ship. Emerging infectious diseases 11, 154 (2005).

11. K. A. Ward, P. Armstrong, J. M. McAnulty, J. M. Iwasenko, D. E. Dwyer, Outbreaks of pandemic (H1N1) 2009 and seasonal influenza A (H3N2) on cruise ship. Emerging infectious diseases 16, 1731 (2010).

12. J. D. Malone, USS Theodore Roosevelt, COVID-19, and ships: lessons learned. JAMA network open 3, e2022095–e2022095 (2020).

13. L. F. Moriarty, Public health responses to COVID-19 outbreaks on cruise ships—worldwide, February–March 2020. MMWR. Morbidity and mortality weekly report 69, (2020).

14. L. Huang et al., Rapid asymptomatic transmission of COVID-19 during the incubation period demonstrating strong infectivity in a cluster of youngsters aged 16-23 years outside Wuhan and characteristics of young patients with COVID-19: a prospective contact-tracing study. Journal of Infection, (2020).

15. A. Sakurai et al., Natural History of Asymptomatic SARS-CoV-2 Infection. New England Journal of Medicine, (2020).

16. A. J. Hall, M. E. Wikswo, K. Pringle, L. H. Gould, U. D. Parashar, Vital signs: foodborne norovirus outbreaks—United States, 2009–2012. MMWR. Morbidity and mortality weekly report 63, 491 (2014).

17. H. Heesterbeek et al., Modeling infectious disease dynamics in the complex landscape of global health. Science 347, aaa4339 (2015).

18. E. Kajita, J. T. Okano, E. N. Bodine, S. P. Layne, S. Blower, Modelling an outbreak of an emerging pathogen. Nature Reviews Microbiology 5, 700–709 (2007).

19. S. Eubank et al., Modelling disease outbreaks in realistic urban social networks. Nature 429, 180–184 (2004).

20. A. J. Kucharski et al., Early dynamics of transmission and control of COVID-19: a mathematical modelling study. The lancet infectious diseases 20, 553–558 (2020).

21. H. Arduin, M. Domenech de Cellès, D. Guillemot, L. Watier, L. Opatowski, An agent-based model simulation of influenza interactions at the host level: insight into the influenza-related burden of pneumococcal infections. BMC infectious diseases 17, 1–12 (2017).

22. M. Tracy, M. Cerdá, K. M. Keyes, Agent-based modeling in public health: current applications and future directions. Annual review of public health 39, 77–94 (2018).

23. B. Faucher et al., Agent-based modelling of reactive vaccination of workplaces and schools against COVID-19. Nature communications 13, 1414 (2022).

24. A. S. Salehi, P. Garner, Occupational injury history and universal precautions awareness: a survey in Kabul hospital staff. BMC infectious diseases 10, 1–4 (2010).

25. R. Cassidy et al., Mathematical modelling for health systems research: a systematic review of system dynamics and agent-based models. BMC health services research 19, 1–24 (2019).

26. E. Hunter, B. Mac Namee, J. Kelleher, An open-data-driven agent-based model to simulate infectious disease outbreaks. PloS one 13, e0208775 (2018).

27. R. Hinch et al., OpenABM-Covid19—An agent-based model for non-pharmaceutical interventions against COVID-19 including contact tracing. PLoS computational biology 17, e1009146 (2021).

28. N. Hoertel et al., A stochastic agent-based model of the SARS-CoV-2 epidemic in France. Nature medicine 26, 1417–1421 (2020).

29. V. Thomopoulos, K. Tsichlas, An agent-based model for disease epidemics in greece. Information 15, 150 (2024).

30. R. J. Rockett et al., Revealing COVID-19 transmission in Australia by SARS-CoV-2 genome sequencing and agent-based modeling. Nature medicine 26, 1398–1404 (2020).

31. G. Rutherford, M. R. Friesen, R. D. McLeod, An agent based model for simulating the spread of sexually transmitted infections. Online Journal of Public Health Informatics 4, (2012).

32. L. Perez, S. Dragicevic, An agent-based approach for modeling dynamics of contagious disease spread. International journal of health geographics 8, 1–17 (2009).

33. W. O. Kermack, A. G. McKendrick, A contribution to the mathematical theory of epidemics. Proceedings of the royal society of london. Series A, Containing papers of a mathematical and physical character 115, 700–721 (1927).

34. M. Y. Li, J. S. Muldowney, Global stability for the SEIR model in epidemiology. Mathematical biosciences 125, 155–164 (1995).

35. G. Wilmink et al., Real-time digital contact tracing: development of a system to control COVID-19 outbreaks in nursing homes and long-term care facilities. JMIR public health and surveillance 6, e20828 (2020).

36. G. J. Smith et al., Origins and evolutionary genomics of the 2009 swine-origin H1N1 influenza A epidemic. Nature 459, 1122–1125 (2009).

37. C. Wang, P. W. Horby, F. G. Hayden, G. F. Gao, A novel coronavirus outbreak of global health concern. The lancet 395, 470–473 (2020).

38. S. Srinivasan et al., Structural genomics of SARS-CoV-2 indicates evolutionary conserved functional regions of viral proteins. Viruses 12, 360 (2020).

39. S. Baize et al., Emergence of Zaire Ebola virus disease in Guinea. New England Journal of Medicine 371, 1418–1425 (2014).

40. V. J. Munster et al., Pathogenesis and transmission of swine-origin 2009 A (H1N1) influenza virus in ferrets. Science 325, 481–483 (2009).

41. G. P. Kobinger et al., Replication, pathogenicity, shedding, and transmission of Zaire ebolavirus in pigs. Journal of Infectious Diseases 204, 200–208 (2011).

42. R. L. Atmar et al., Norwalk virus shedding after experimental human infection. Emerging infectious diseases 14, 1553 (2008).

43. K. K.-W. To et al., Consistent detection of 2019 novel coronavirus in saliva. Clinical Infectious Diseases, (2020).

44. D. K. Ip et al., Viral shedding and transmission potential of asymptomatic and paucisymptomatic influenza virus infections in the community. Clinical infectious diseases 64, 736–742 (2017).

45. J. Papenburg et al., Household transmission of the 2009 pandemic A/H1N1 influenza virus: elevated laboratory-confirmed secondary attack rates and evidence of asymptomatic infections. Clinical Infectious Diseases 51, 1033–1041 (2010).

46. P. Teunis et al., Shedding of norovirus in symptomatic and asymptomatic infections. Epidemiology & Infection 143, 1710–1717 (2015).

47. P. Mlcochova et al., SARS-CoV-2 B. 1.617. 2 Delta variant replication and immune evasion. Nature, 1–6 (2021).

48. I. F.-N. Hung et al., SARS-CoV-2 shedding and seroconversion among passengers quarantined after disembarking a cruise ship: a case series. The Lancet Infectious Diseases, (2020).

49. A. N. Kraay et al., Fomite-mediated transmission as a sufficient pathway: a comparative analysis across three viral pathogens. BMC infectious diseases 18, 1–13 (2018).

50. CDC.

51. P. F. Teunis, N. J. Nagelkerke, C. N. Haas, Dose response models for infectious gastroenteritis. Risk Analysis 19, 1251–1260 (1999).

52. P. Teunis, A. Havelaar, The beta Poisson dose-response model is not a single-hit model. Risk Analysis 20, 513–520 (2000).

53. P. F. Teunis et al., Norwalk virus: how infectious is it? Journal of medical virology 80, 1468–1476 (2008).

54. S. Luke, C. Cioffi-Revilla, L. Panait, K. Sullivan, G. Balan, Mason: A multiagent simulation environment. Simulation 81, 517–527 (2005).

55. T. Ormsby, Getting to know ArcGIS desktop: basics of ArcView, ArcEditor, and ArcInfo. (ESRI, Inc., 2004).

56. M. Loy, R. Eckstein, D. Wood, J. Elliott, B. Cole, Java swing. (“O’Reilly Media, Inc.”, 2002).

57. C. J. Willmott, K. Matsuura, Advantages of the mean absolute error (MAE) over the root mean square error (RMSE) in assessing average model performance. Climate research 30, 79–82 (2005).

58. F. J. Massey Jr, The Kolmogorov-Smirnov test for goodness of fit. Journal of the American statistical Association 46, 68–78 (1951).

59. M. W. Fong et al., Nonpharmaceutical measures for pandemic influenza in nonhealthcare settings—social distancing measures. Emerging infectious diseases 26, 976 (2020).

60. A. Q. Ohi, M. Mridha, M. M. Monowar, M. A. Hamid, Exploring optimal control of epidemic spread using reinforcement learning. Scientific reports 10, 22106 (2020).

61. Y. Liu et al., Aerodynamic analysis of SARS-CoV-2 in two Wuhan hospitals. Nature 582, 557–560 (2020).

62. Y. Liu, W. Mao, N. Diaz-Elsayed, An investigation of the indoor environment and its influence on manufacturing applications via computational fluid dynamics simulation. Building and Environment 219, 109161 (2022).

63. C. M. Brown, Outbreak of SARS-CoV-2 infections, including COVID-19 vaccine breakthrough infections, associated with large public gatherings—Barnstable County, Massachusetts, July 2021. MMWR. Morbidity and mortality weekly report 70, (2021).

