## Supplementary Information for "Real-Time Spatiotemporal Tracking of Infectious Outbreaks in Confined Environments with a Host-Pathogen Agent-Based System"

*This page intentionally left blank*

### Supplementary Methods

#### 1. Additional details on AI-GIS Infection Dynamics (AGID) model

The entire approach or modeling system architecture includes 4 basic components: (1) Vessel representation and real-time dashboard; (2) Population representation and action modeling; (3) Pathogen model; and (4) Containment protocols.

##### 1.1. Vessel representation and real-time dashboard

A 3D representation of a specific cruise ship's (Ship X) floor plan was rendered in a geographical information system (GIS) (Fig. 1A). To accomplish this, images of the floor plan for each level of the cruise ship were obtained from a publicly available source. These raster images included details on how the levels of the ship are accessible via stairs and elevators. Next, the floor plan in each image was vectorized using Inkscape (https://inkscape.org). Given the initial vectorizations, a polygonal representation of the areas on each level of the ship (*e.g.*, staterooms, restaurants, hallways, etc.) was created using ArcGIS (https://www.arcgis.com/), a geographic information software package. The polygons for the spaces on each level were then partitioned into smaller areas representing locations that agents could inhabit at any point in time. Finally, the locations (nodes) were connected using paths (edges) corresponding to the adjacency of these locations as present in the floor plans, which created a traversable network. The completed network model was then used in Mason, an open source multiagent simulation library (*1*), as the environment within which the movements of onboard members can be modeled. The simulation is primarily driven by an internal clock that is represented by a step-counter based on which all the actions in the simulated world take place. Each step equals one second in the simulated world, *e.g.*, one simulated hour = 3,600 steps. Through this internal clock we create the desired length of the cruise to match a specific number of days and it drives the agent behavior on an hourly and daily basis. The simulation framework is a single-threaded process and because of the internal clock, the speed of the simulation is directly proportional to the single-thread clock speed of the processor, *e.g.*, the simulation of a 10-day cruise takes less than 4 hours on an Intel Core i7-6700K processor that has single and multicore clock speeds of 4 GHz. The number of simultaneous simulations scales linearly based on the number of available physical CPU cores.

The real-time dashboard has two windows for interacting with the system (Fig. 1B), a primary graphical user interface (GUI) containing the GIS, agents and real-time infection statistics and a secondary GUI to start, pause, stop and control the speed of the simulation. The major components of the GUI are rendered using Java Swing (*2*), and the agents and statistics are rendered using Mason. In the primary GUI, there are five informational areas (Fig. 2A), the largest of them is the GIS with the live agents. On the right side of the GIS is the legend indicating the color coding for the various areas on the cruise ship. The legend also includes color codes for the infection phases in passenger and crew and a notification icon which appears in the location where an agent became ill. Below the GIS are two plots, the left plot displays real-time infection statistics as a function of the number of steps in the simulation, and the right plot displays the daily infection cases. The final informational area in the GUI is in the bottom right corner, where a magnified view of the GIS and agents is available, to provide a detailed view of high-activity areas such as dining rooms. In addition to the informational areas in the primary GUI, there are some utilities available to the user, such as functionality to capture a snapshot of the dashboard, specifying the magnification of high-activity area and both horizontal and vertical scrolling of the GIS.

In addition to the GUI-based dashboard that shows two summary statistics of infection, we also implemented a reporting system that logs detailed infection statistics in the background to a file. The infection statistics are reported separately for passenger and crew and included statistics such as: healthy, infected, symptomatic, asymptomatic, recovered and dead. The data logging starts at 6 AM on the first day of the cruise and continues at 15-minute intervals (*i.e.*, based on internal clock) till the end of the simulation.

#### 1.2. Population representation and action modeling

To accurately model the population in Ship X, we first find the number of passengers and number of crew onboard from information available to the public. The maximum onboard capacity of Ship X is 2,702 persons of which approximately 70% (1,888) are passengers and 30% are crew (814). We model the daily activity for passengers and crew (Fig. 1), where the agents spend a certain amount of time in an area of the ship while they perform the activity. Using the GIS network and the defined activity, an agent traverses the ship and goes to a particular location to perform the necessary activity *e.g.*, going from their stateroom to a dining area for breakfast and staying there until end of meal.

The passengers and crew are modeled as two separate types of agents because of their daily schedules (Supp. Fig. S10 and S11). The daily schedules of the passengers are modeled with stochasticity as observed in the real-world *i.e.*, the leisure activity performed by the agent is selected randomly, such that not all agents are performing the same activity at any given time. To resemble activities taking place on cruise ships, some events such as mealtimes are important since they have 2-3 hours of time window during which people visit the dining areas and spend a significant amount of time, such schedules create a large group of agents in a single location. For other activities such as leisure time, there is no strict timeframe and the agents traverse various recreational areas and spend an arbitrary amount of time.

#### 1.3. Pathogen model

The pathogen model incorporates all the infection related attributes of a disease-causing organism to effectively simulate an outbreak and specifically in our investigations, viral outbreaks. Specifically, we integrate two complementary paradigms of infection, (i) dose-response relationship and (ii) hit theory of infection. We model all the major phases of an infection (Supp. Fig. S13), *i.e.*, exposure, infection, illness and the outcomes of either recovery/sequelae/death. Each virus has a set of important attributes/parameters which determine how serious the outbreak could be, these are provided (Supp. Table S4).

In the first paradigm of infection, the dose-response relationship (*3*) aims to study the outcome in a host when exposed to a certain quantity of a pathogenic microorganism. Hence, one of the most important parameters is the viral shedding value (Supp. Table S4). These values represent the number of viral particles shed by the infected host obtained through titers of sampled sites *e.g.*, oropharynx (Fig. 1 and Supp. Fig. S6). Information on viral shedding is important because an infected host can shed viral particles when symptomatically and asymptomatically (*i.e.*, in the prodromal period and/or during convalescence) (*4*). Although the number of viral particles are generally fewer during asymptomatic shedding, it still creates a favorable condition for disease transmission as the host and persons in close contact are unaware of the infection and hence cannot take preventative measures to reduce transmission (*5*).

Since the modes of transmission vary for each pathogen (*6*), *i.e.*, aerosol-based and droplet-based. the viral shedding values obtained in laboratories from samples (*7*) are higher than the viral particles that are transmitted through various modes (*8*). Thus, we also define a dose adjustment parameter to control the actual quantity of contaminated material a healthy agent will be exposed to, based on the shedding value of an infected host. For instance, in a simulation, an agent shedding $10^{11}$ viral particles per gram would expose a single healthy individual to a dose $10^{6}$-$10^{3}$ copies (a dose from this range is assigned randomly).

In the second paradigm, the hit theory of infection (*9*) defines the pathogenic infection of a host as the outcome of a sequence of events. For this purpose, our pathogen model works in conjunction with our agent model to completely simulate an infection, *i.e.*, the agents can get the disease and act as vectors to transmit the disease. Depending on the type of virus and on being infected, an agent generally can go through the following four events: (1) exposure, (2) infection, (3) illness and (4) recovery/sequelae/death. Of these, there are two important sequential events: exposure to certain quantity of viral particles which can lead to infection, and infection that can lead to illness, which we mathematically define as a probability (*9*):

$$\mathrm{P(ill|dose)} = \mathrm{P(ill|inf)} * \mathrm{P(inf|dose)}.$$

In the above equation, the probability of being infected when exposed to a specific quantity of viral particles is defined by the equation:

$$\mathrm{P(inf|dose)} = \mathrm{P_{inf}}(\mathrm{D}; \alpha, \beta) = 1 - (1 + \frac{\mathrm{D}}{\beta})^{-\alpha},$$

where D is the dose of viral particles, and α and β are infectivity parameters obtained from literature (*10, 11*).

If a host becomes infected after exposure, then the incubation period is initialized according to the virus being modeled. During the incubation period is when the agent can develop the illness, which we define as the probability of progressing to illness given by the following equation:

$$\mathrm{h}(\mathrm{D}|\eta, \mathrm{r}) = 1 - (1 + \eta\mathrm{D})^{-\mathrm{r}},$$

where D is the dose of viral particles, and η and r are maximum-likelihood estimates obtained from literature (*10, 11*).

If an agent does not get ill from the viral particles that infected it, then that agent progresses directly to the recovery phase. However, if an agent progresses to the illness phase, we change certain behaviors to match what is observed in ill people. We reduce the activity levels of the agent by 50% and they spend the remaining time in their staterooms. The period of illness prior to recovery or sequelae or death is specific to the virus being modeled. The important change that occurs in the agent is the number of viral particles they shed during exposed and recovery (asymptomatic shedding) and during illness phase (symptomatic shedding). This change is modeled based on shedding values of each virus, and the result is that an illness phase is when an agent is the most infectious. For certain viruses with a high mortality rate, *e.g.*, Ebola, we model the literature reported fatality rate in the cruise ship population. In this scenario, instead of recovery, a severely ill agent dies from the pathogen and that agent is removed from the simulation to reduce further infection, as observed in the real-world (*12*).

At the end of the period of simulation, the onboard population consists of a certain number of healthy, infected, ill and recovered (dead where appropriate) agents. Most of the cruises (~80%) are less than 10 days long with the most popular (~50%) length of cruise being 6-8 days as reported by CLIA (*13*). Thus, to study the outbreak of an infection extensively and to learn about the outcome where the cruise duration is long *i.e.*, could result in a larger outbreak, we choose to the simulate outbreaks for a period of 10 days. The longer simulation period is also beneficial for shorter cruises due to the real-time nature of the simulation during which the necessary information can be extracted for any day and time.

We selected four viruses to model independent outbreaks on the cruise ship resulting in gastrointestinal illnesses (Norovirus), influenza-like illnesses (H1N1 2009 pandemic virus and COVID-19 virus) and hemorrhagic fever (Zaire ebolavirus). The viruses were selected such that we could model variation in transmission modes, disease cycles, attack rates fatality rates and the impact of introduction of novel pathogens. The shedding values for H1N1 2009 pandemic strain (H1N1pdm09) and ebolavirus (EBOV) were obtained from animal model studies (*14, 15*) norovirus from human challenge studies (*7*) and COVID-19 virus (SARS-CoV-2) from clinical testing (*16*). A summary of the characteristics of the viruses is provided including the shedding value or dose-response curves (Supp. Table S2 and Supp. Fig. S5).

Since SARS-CoV-2 was an emerging pandemic there has been a well-documented outbreak at the beginning aboard the cruise ship Diamond Princess of the coast of Japan. To understand the dynamics of that specific outbreak we recreate the environment since Diamond Princess was similar in size and onboard capacity as Cruise Ship X. In addition, we model a few scenarios that could have resulted in the observed outbreak and temporal dynamics. Specifically, we model different incubation periods and asymptomatic transmission rates that were noted on the ship and other facilities such as nursing homes (*17, 18*) including fatality rates noted in the early months of the pandemic (*19, 20*).

For airborne pathogens like SARS-CoV-2, the asymptomatic transmission plays a crucial role in the transmission of the disease (*21*) and similar dynamics were observed during the 2009 H1N1 pandemic as well (*22*) where the virus shedding is significant in asymptomatic cases. This is unlike pathogens that spread through direct contact such as Norovirus and EBOV where asymptomatic cases do not shed sufficient pathogens to transmit the disease (*4, 23*). In addition to the wild type SARS-CoV-2 virus, we also modeled the Delta variant which has a smaller incubation period, higher viral load and higher rates of transmission (*24*).

##### 1.4. Containment protocols

We model various containment protocols to evaluate strategies that can limit the spread of an infection. Containment strategies/protocols are necessary to modify community and individual behavior and reduce the spread of an infectious outbreak to ultimately "flatten the curve" (*25*). Containment strategies are designed based on the environment where an outbreak occurs and are established by a governing agency such as the United States Centers for Disease Control and Prevention (CDC). For the cruise ship industry, the US CDC has specifically established the vessel sanitation program (VSP) to prevent and control gastrointestinal outbreaks onboard a ship (*26*). The VSP includes a detailed operations manual that is updated approximately every 5 years, crew training and ship inspections, and surveys have shown that inspection scores have gradually improved over time (*27*). The directive for VSP to be initiated is when 3% of passengers or crew report with symptoms of gastrointestinal illness (GII).

However, the CDC does not have a similarly rigorous program for influenza-like illnesses (ILI) (*26*). Cruise ships are expected to only report ILI if the cases are equal to or greater than 1.38 cases per 1,000 passenger-days (*28*). This is given by the following equation, $n = (1.38)(p \times d) \div 1000$, where $p$ is total number of passengers or crew, $d$ is number of days on the voyage at time of reporting to CDC quarantine station. In addition, the passengers and crew are recommended to follow general guidelines, in a case-by-case manner, to control the spread of ILI. However, these general guidelines might not be sufficient for unusual ILIs, as observed with the novel coronavirus and more stringent, ship-wide measures might be required to control the spread.

Therefore, to contain the spread of GII and ILI we implemented four containment protocols (Supp. Table S3). According to the Protocol-1, on identifying a food-borne cause for the GII, we close only the dining area where the infection started for thorough cleaning and isolate the affected persons. The rationale for the Protocol-2 is to shut down all dining areas as they are the highest density areas on a ship. Once the outbreak has started, this can significantly reduce transmission since persons shedding the virus will not come in contact with people in dining areas. The symptomatic persons are isolated and all meals are delivered to the staterooms. The first two protocols are designed with consideration for GIIs.

Protocol-3 is a modified version of the CDC VSP, where we incrementally reduce the case threshold parameter (*i.e.*, CDC recommended 3% of passengers/crew). This protocol implements the CDC validated cleaning and isolation measures but with a more stringent threshold to further reduce the outbreak size and observe the outcome w.r.t GIIs and ILIs.

The final protocol, Protocol-4 is a classic social distancing measure, where the individuals who develop any symptom immediately isolate themselves in their room and continue the isolation until 48-hours after all their symptoms are cleared without the need for medications. Protocol-4 is more stringent than ILI guidelines from CDC, as they only recommend isolating until 24-hours after the resolution of fever and do not appreciably consider other symptoms through which viral shedding can occur in the convalescence phase, a coughing etiquette is recommended.

SARS-CoV-2 is an emerging infection with varying symptomatology that can mimic a respiratory or gastrointestinal illness among a myriad set of symptoms (*29, 30*). As the CDC guidelines for GIIs are more stringent than that for ILIs. For the wild type SARS-CoV-2 infections, in addition to the default VSP threshold of 3%, we implemented stricter thresholds of the CDC VSP at 2% and 1% to study the effectiveness in outbreak control. For the SARS-CoV-2 Delta variant, we implemented face masks with data from recent studies on SARS-CoV-2 community transmission (*31*) and performed simulations to study the effect on case load without prohibiting any activities.

#### 2. Evaluation protocol

To assess the predictions from the various pathogen and containment modeling, we design a rigorous simulation and evaluation protocol. The details are provided below.

##### 2.1. Simulation criterion for outbreaks

To have sufficient predictive capability while also maintaining computational feasibility, specific number of simulations are performed for each outbreak model for cruise ship X to obtain the average values of all the statistics. Specifically, the combination of each pathogen and containment protocol is simulated ten times to obtain summary infection statistics. To compute the daily infection averages across all the ten simulations for each setting, we consistently sample the data at 7 AM on each day of the cruise. As a crucial criterion, any parameter of the system that is modified to study the outcome is simulated ten times. Thus, for each pathogen and for each protocol we have a total of 160 simulations for Cruise Ship X. As part of validation, we have eight types of naïve simulations that contribute an additional 80 simulations.

#### 2.2. Verifying the modeling capability with real outbreaks

For a system that models outbreaks, it is necessary to verify the generalizability of that system under various modeling requirements. Since the CDC maintains historical data about norovirus outbreaks on cruise ships, we can use this data to verify the generalizability of our system to model outbreaks on various cruise ships.

We would like to test the system by modeling a specific real outbreak that occurred on Cruise Ship X. For Cruise Ship X, we were able to collect detailed per-day data about an outbreak from 2006 that happened over a nine-day period. Using some of the collected data to seed the initial parameters of the system, we compare how accurate the outcome of the simulations is to those of the real outbreak. We use only the initial number of ill members to seed the disease vectors and perform ten simulations.

First, to show the benefits and accuracy of detailed agent and pathogen modeling, we create a few indiscriminate models. There are three types of naïve simulations based on: (1) naïve agent behavior and formal pathogen characteristics, (2) naïve agent behavior and naïve pathogen characteristics and (3) formal agent behavior and naïve pathogen characteristics. Specifically, we create eight different models where the agent and pathogen models are lacking detailed parameters and are replaced with some naïve/random parameters. To begin with Type-1, in naïve simulation-1 (NS-1), the agents do not have any scheduling and traverse the ship throughout the day without any breaks/sleep and in naïve simulation-2, the agents still perform a randomized traversal of the ship, except they have breakfast in the dining areas. Naïve simulations-3, 4 & 5 are part of Type-2, where the agents still perform a randomized traversal of the ship, except now the pathogen characteristics are changed. In NS-3: the agents are infectious all the time; NS-4: an infectious agent transmits the pathogen to all agents in the vicinity and in NS-5: the agents are infectious all the time and infect everyone in the vicinity. As part of Type-3, naïve simulations-6, 7 & 8 follow the pathogen characteristics as detailed in NS-3, 4 & 5 respectively, but the agents follow a detailed schedule of a typical day aboard the ship.

Second, to systematically compare the predictions of the models with the real outbreak, we use multiple metrics that are detailed below. The first metric, root-mean-square error (RMSE), is commonly used to quantify the difference between observed values and predicted values in forecasting scenarios (which fits our goal to forecast an outbreak). Root-mean-square deviation is defined as,

$$RMSE = \frac{\sqrt{\sum_{i=1}^{n}(y_i - \hat{y_i})^2}}{n}$$

where $n$ is the number of expected/predicted values, $y_i$ is the observed value and $\hat{y_i}$ is the predicted value. The interpretation is $0 \leq RMSE \leq \infty$ , therefore the ideal model has a $RMSE = 0$, *i.e.*, all predicted values match observed values. As an auxiliary model evaluation metric to better quantify average model error, we also use mean absolute error (MAE). As RMSE assigns a larger weight to errors of higher magnitude (*32*), MAE provides a linear measure of errors.

$$MAE = \frac{\sum_{i=1}^{n}|y_i - \hat{y_i}|}{n}$$

Since the predicted number of daily cases can be considered as resulting from a probability distribution function (PDF), we also consider a second metric, Kolmogorov-Smirnov (K-S) test. The K-S test is a non-parametric test and measure of the distance between a reference probability distribution and a sample probability distribution. Specifically, we use the two-sample K-S test as a "goodness of fit" test (*33*), to compare the cases of the simulated outbreak with the real outbreak cases. KS test is generally defined as

$$D_{m,n} = \max_x \left| F_{1,m}(x) - F_{2,n}(x) \right|$$

Where $F_{1,m}$ and $F_{2,n}$ are the empirical distribution functions for $m$ and $n$ samples, respectively, in our case $m = n$. The test statistic $D$ represents the maximum distance between the empirical distribution functions or their curves *i.e.*, smaller value represents higher similarity. The null hypothesis, $H_0$, is that both samples are drawn from the same distribution and the $H_0$ is rejected at significance level $\alpha$ if

$$D > \frac{1}{\sqrt{n}} \sqrt{-log\left(\frac{\alpha}{2}\right)}$$

Additionally, we compare the time taken to reach a case load of 3% of the population *i.e.*, the critical threshold for CDC VSP. This is necessary to quantify the rate at which the simulated infection grows with respect to the growth rate observed in the real outbreak. In addition to the per-day forecast, we summarize the infections through attack rate *i.e.*, percentage of population at risk that contracts the infection for a pathogen and under a containment scenario. To plan mitigations, it is necessary to know in advance when the peak of an infection can occur (*34*). To facilitate this, we convert the aggregated per day infection data to probability densities to provide the likelihood for infection peaks within a range of days. We also analyze the shift in the probability of peaks under the different containment protocols.

Third, we would like to compare the simulated outbreak and the real outbreak to a mathematical modeling of a norovirus outbreak on cruise ship X. For this purpose, in the fixed population we use the widely applied compartmental model of susceptible (S), infectious (I) and removed (R) individuals (*35*). The entire model is a function of time $t$, and the dependent variables are S, I and R. The SIR model is represented by the following set of ordinary differential equations.

$$\frac{dS}{dt} = -\beta SI$$

$$\frac{dI}{dt} = \beta SI - \gamma I$$

$$\frac{dR}{dt} = \gamma I$$

Where $N$ is the constant number of individuals in the population (*i.e.*, no birth or death), such that $N = S + I + R$. $\gamma$ is the average period of infectiousness and $\beta$ is the transmission rate $\beta = R_0\gamma$, using which the basic reproduction number is defined as $R_0 = \frac{\beta}{\gamma}$. There is no incubation period in the SIR model.

We also apply the SEIR compartmental model, which has the four compartments of Susceptible – Exposed – Infectious – Recovered respectively. The additional Exposed compartment models the population that is infected but not yet infectious, to account for the incubation period defined by $\sigma$. The SEIR model is defined by the following equations, where all other terms are shared with the SIR model.

$$\frac{dS}{dt} = -\beta SI$$

$$\frac{dE}{dt} = \beta SI - \sigma I$$

$$\frac{dI}{dt} = \sigma E - \gamma I$$

$$\frac{dR}{dt} = -\gamma I$$

Fourth, the above three evaluation criteria were based on the temporal dynamics of the simulated outbreak, hence we also plan to compare the spatial dynamics of the transmissions during the outbreak. For this purpose, we collect the coordinates of the infections from the GIS and create a location probability density and overlay it on the ship areas as a heatmap. This provides an intuitive visualization through which infection hotspots on the ship can be identified and compared to recorded data in the CDC outbreak reports. We also study the changes that occur in the infection hotspots under the various containment protocols.

So far, we have assessed if the proposed system can accurately simulate the outbreak for a particular pathogen on a specific cruise ship. After satisfying the previous criteria, we can now evaluate the generalizability of our system for the same pathogen but in cruise ships of various physical sizes and population sizes by following a rigorous procedure. To reuse the GIS of Cruise Ship X, we carefully collect data on many cruise ships ranging from similarly large to small ships *i.e.*, they have the same or fewer number of areas (staterooms, dining rooms etc.) and a proportional onboard population. To achieve this, we collected data on five cruise ships (Supp. Table S1). To obtain data about the range of outbreak size in each ship, a ship was selected such that at least five norovirus outbreaks had occurred. This summary data collected from CDC serves the purpose of comparing the general trends in simulated outbreaks from many cruise ships. For each cruise ship, we simulate norovirus outbreaks under the CDC VSP scenario.

By benchmarking the system in various cruise ships against norovirus outbreaks for which data exists, we can test the system's ability to model outbreaks of other pathogens such as EBOV and H1N1pdm09, for which no detailed data exists. However, the single well documented case of COVID-19 outbreak on the Diamond Princess serves us a comparable data source to test the SARS-CoV-2 models against, since it is similar to Ship X in size and onboard capacity.

### Supplementary Tables

**Supplementary Table S1**. Details about the size of the cruise ships used for evaluation along with the design passenger and crew capacities.

| Cruise Ship | Staterooms | Passengers | Crew |
|---|---:|---:|---:|
| Ship X | 896 | 1,888 | 814 |
| Ship 1 | 630 | 1,258 | 557 |
| Ship 2 | 548 | 1,070 | 655 |
| Ship 3 | 467 | 930 | 465 |
| Ship 4 | 375 | 750 | 542 |
| Ship 5 | 345 | 690 | 408 |

**Supplementary Table S2**. Primary characteristics of the four viruses modeled to cause outbreaks on cruise ships.

| **Virus** | Norovirus | EBOV | H1N1pdm09 | SARS-CoV-2 |
|---|---|---|---|---|
| **Illness** | Gastrointestinal | Hemorrhagic fever | Influenza-like | Acute respiratory distress |
| **Primary mode of transmission** | Contact | Contact | Airborne | Airborne |
| **$R_0$** | 1.6 | 1.9 | 1.3 | 2 |
| **Avg. Incubation period** | 1 – 3 days | 8 – 10 days | 1 – 3 days | 5 – 7 days |
| **Asymptomatic Transmission** | Unlikely | Unlikely | 10% | 50% |
| **Fatality rate** | 0.005% | 80% | 0.03% | 5% |

**Supplementary Table S3.** Containment protocols implemented. The protocols restrict access to specific locations on the ship or change agent behavior or both.

| Index | Protocol | Definition |
|---|---|---|
| 1 | Affected dining area closed | The dining area where the infection was seeded is closed for cleaning |
| 2 | All dining areas closed | All dining areas are closed to reduce transmission, perform cleaning and meals are delivered to rooms |
| 3 | CDC VSP+ | The case threshold parameter for the initiation of VSP is tunable to reduce outbreak size |
| 4 | Self Isolation | Persons conscientiously isolate themselves when symptoms appear and continue isolation until symptom free |

**Supplementary Table S4.** Parameters of viral infection. The values for the below parameters are obtained from literature survey and studies in human cases or animal models.

| **Parameter** | **Definition** |
|---|---|
| $R_0$ (basic reproduction number) | No. of new infections caused by one infected person |
| Incubation period | Time between exposure and onset of symptoms |
| Viral shedding | Viral progeny secreted from a person over the course of the infection. This is further categorized as symptomatic and asymptomatic shedding. |
| Fatality rate | Proportion of deaths in a population during an outbreak |
| Initial vectors | No. of persons who carry and introduce the infection in an environment. |

#### Supplementary Figures

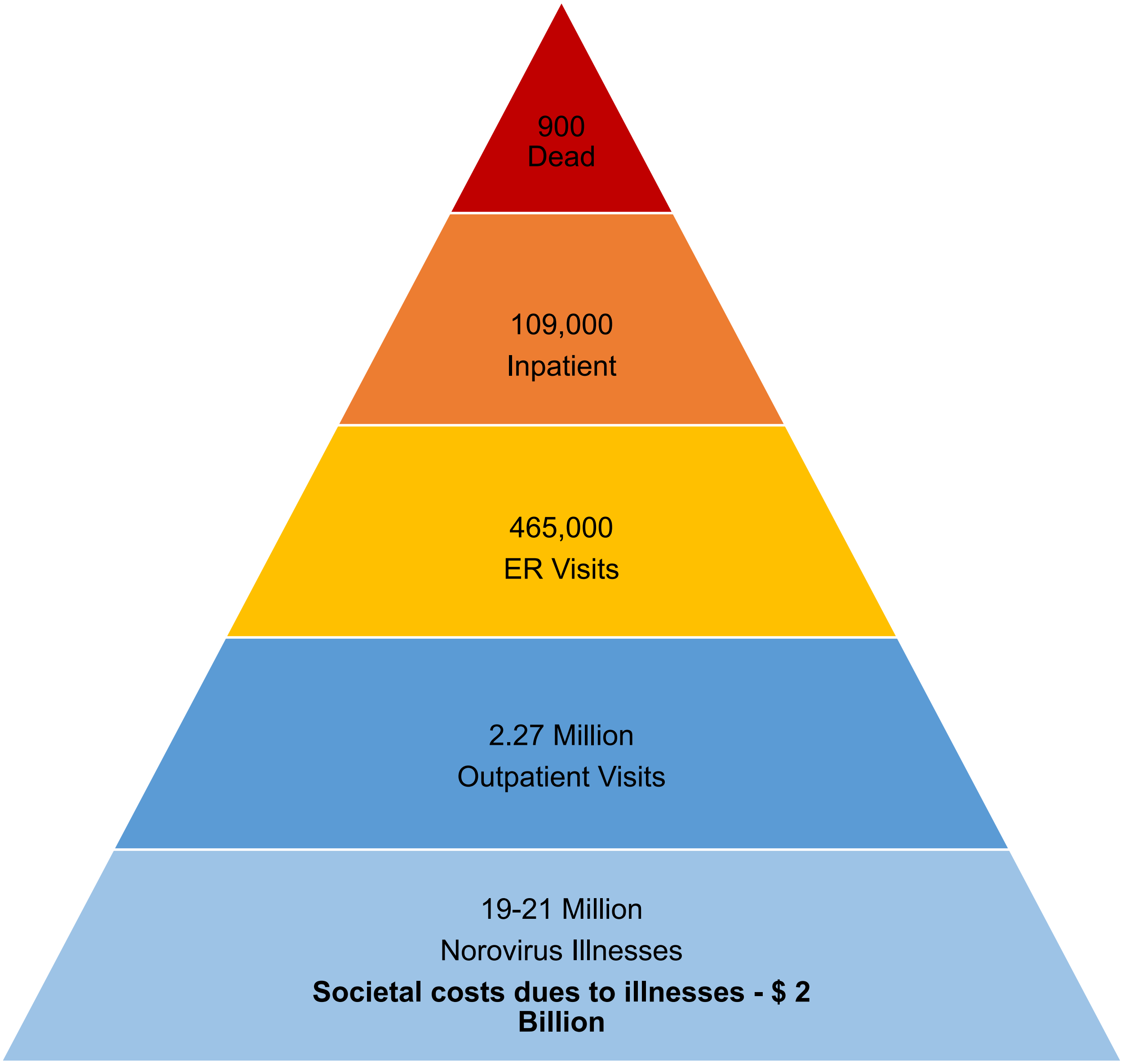


**Supplementary Figure S1.** Norovirus burden in the United States as reported by the Centers for Disease Control and Prevention

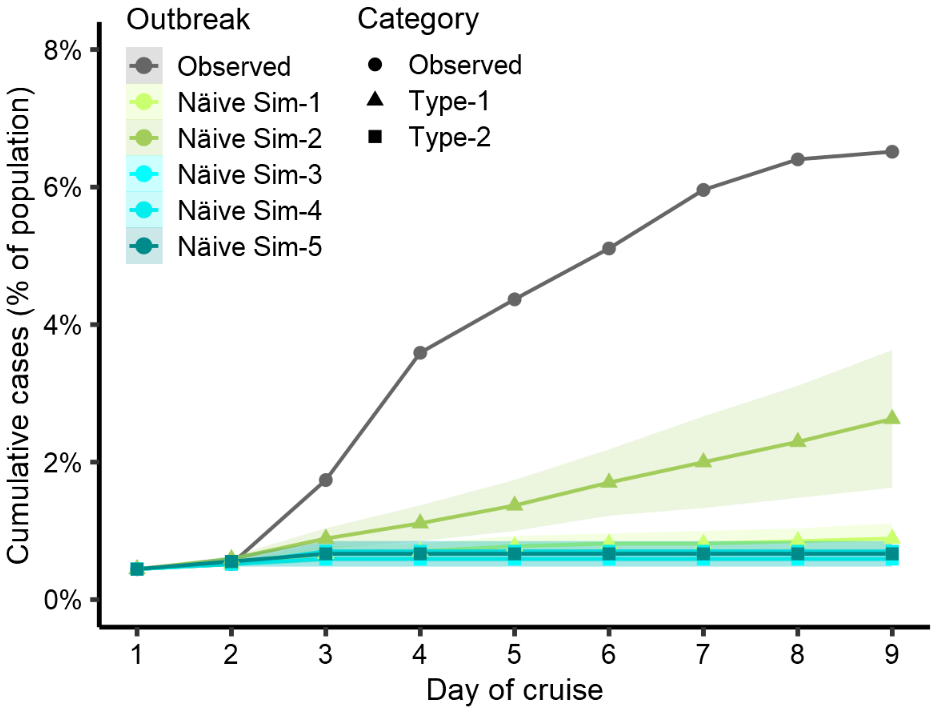


**Supplementary Figure S2.** Naïve agent simulations of Type-1 and 2 with norovirus as the pathogen. In Type-1, for Naïve Sim-1 the agents do not have any scheduling and traverse the ship throughout the day without any breaks/sleep and for naïve sim-2, the agents still perform a randomized traversal of the ship, except they have breakfast in the dining areas. Naïve Sims-3, 4 & 5 are part of Type-2, where the agents still perform a randomized traversal of the ship, except now the pathogen characteristics are changed. In Naïve Sim-3, the agents are infectious all the time; in Naïve Sim-4, an infectious agent transmits the pathogen to all agents in the vicinity; and in Naïve Sim-5, the agents are infectious all the time and infect everyone in the vicinity.

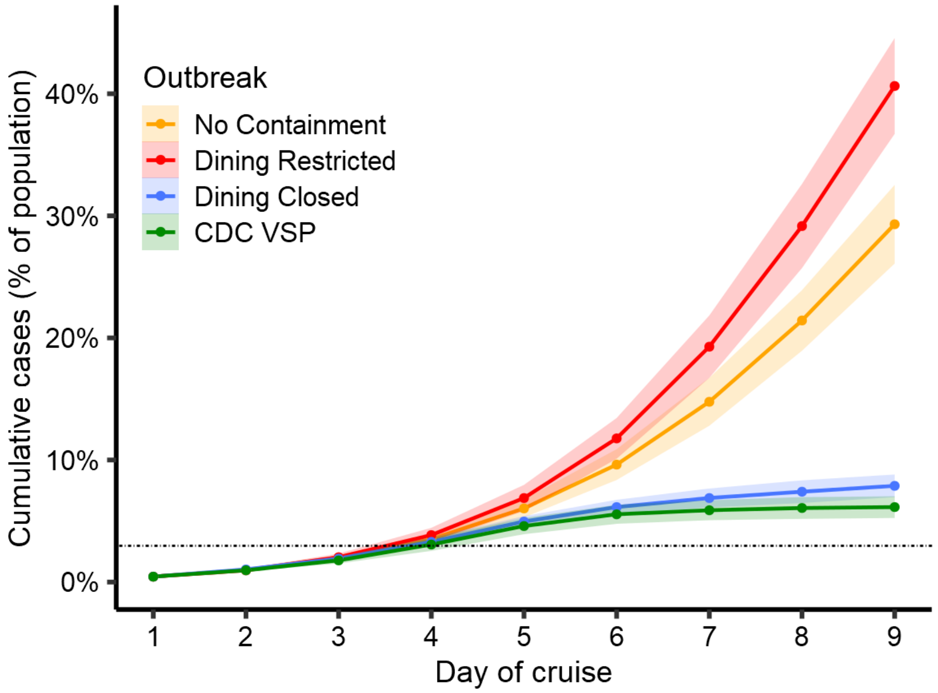


**Supplementary Figure S3.** Additional Norovirus outbreak simulations for Ship X with different containment protocols. The "Dining Restricted" protocol which shuts down a main dining room which is an infection hotspot, results in an increase in case load as it creates crowding in other smaller dining areas. "Dining Closed" protocol which shuts down all dining locations, results in a similar case load as VSP, but with no restriction on other activities.

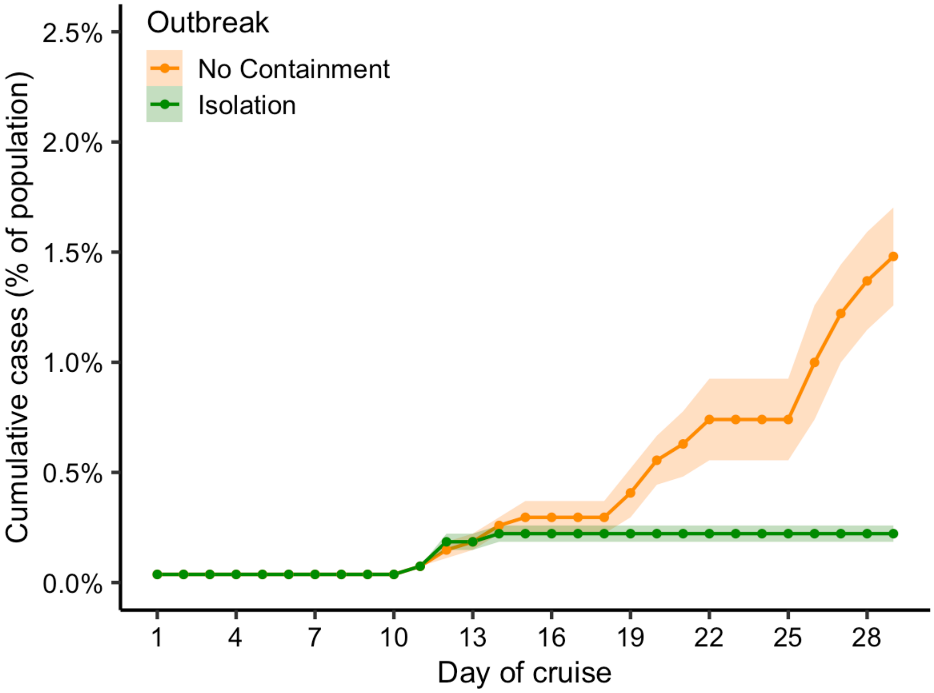


**Supplementary Figure S4.** Simulations for an Ebola outbreak on Ship X. To simulate the Ebola disease lifecycle, which has long incubation periods, the number of days is at the highest duration of typical cruises *i.e.*, 30 days. The long incubation period plays a critical role in a slow start of the outbreak and even with no containment protocols, it has an attack rate of less than 2% by the end of the cruise. Isolation works well for this disease due to the varying long incubation periods, differing symptoms and initial small case load which do not meet the criteria for VSP.

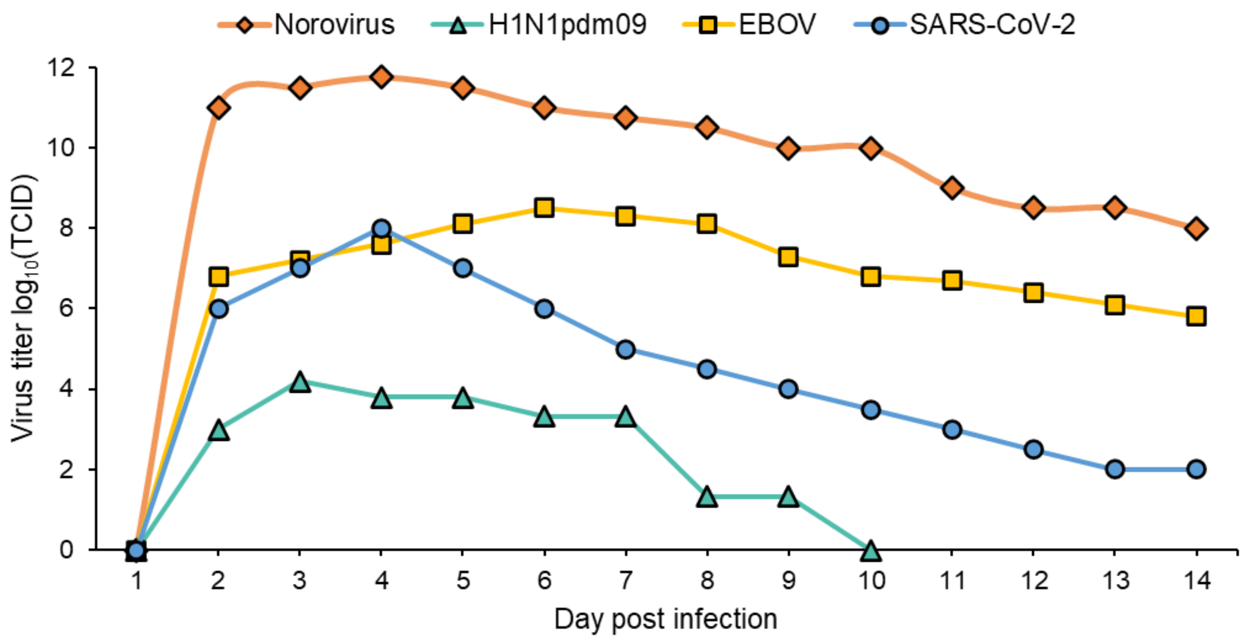


**Supplementary Figure S5.** Shedding values for the four modeled viruses. The shedding values for H1N1pdm09 and EBOV were obtained from animal model studies, norovirus from human challenge studies, and SARS-CoV-2 from clinical testing.

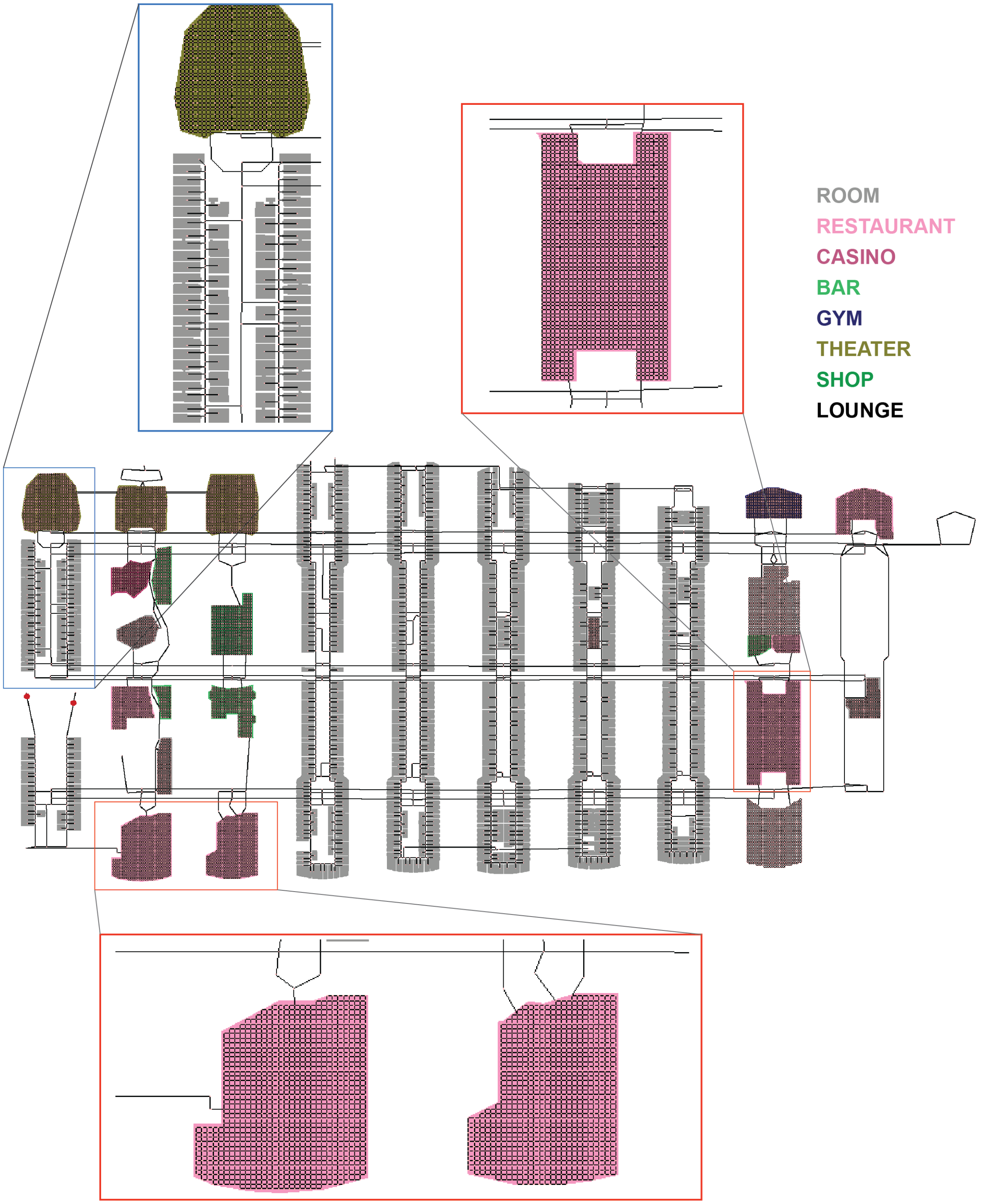


**Supplementary Figure S6.** Color annotation of the main areas of Cruiseship X. Zoomed in are the key areas that appeared as the hotspots in the outbreak simulations caused by different pathogens: dining areas (red squares) and the main crew corridor (blue square).

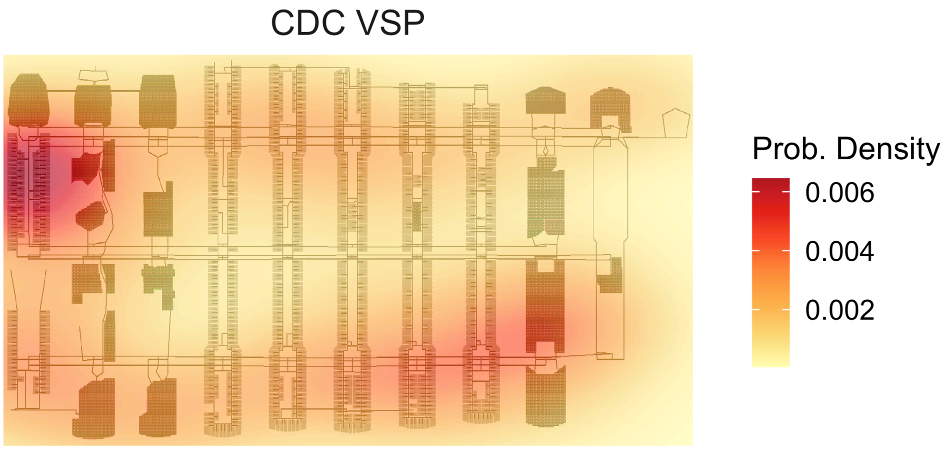


**Supplementary Figure S7.** Geographic distribution of infection cases caused by H1N1pdm09 on Ship X during the 14-day cruise. The infection cases recorded in Fig. 4A show that the Vessel Sanitation Program is not the best strategy (AR: 11%) for a pathogen with primary mode of transmission through air (color scale shows two-dimensional kernel density estimate for hotspots). The proposed Isolation protocol results in only 8 infection cases.

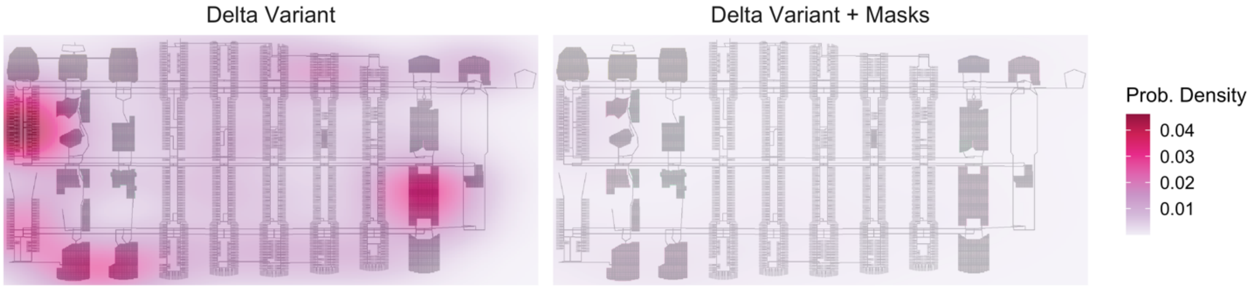


**Supplementary Figure S8.** Geographic distribution of SARS-CoV-2 (Delta variant) infections with different containment protocols. The left panel shows the distribution of infections under the same containment measures that were implemented on the Diamond Princess ship. The attack rate for the Delta variant was 61% compared to 19% for the wild type virus (color scale shows two-dimensional kernel density estimate for hotspots). When masks are implemented for the agents, the attack rate reduces to approx. 7%, shown in the right panel.

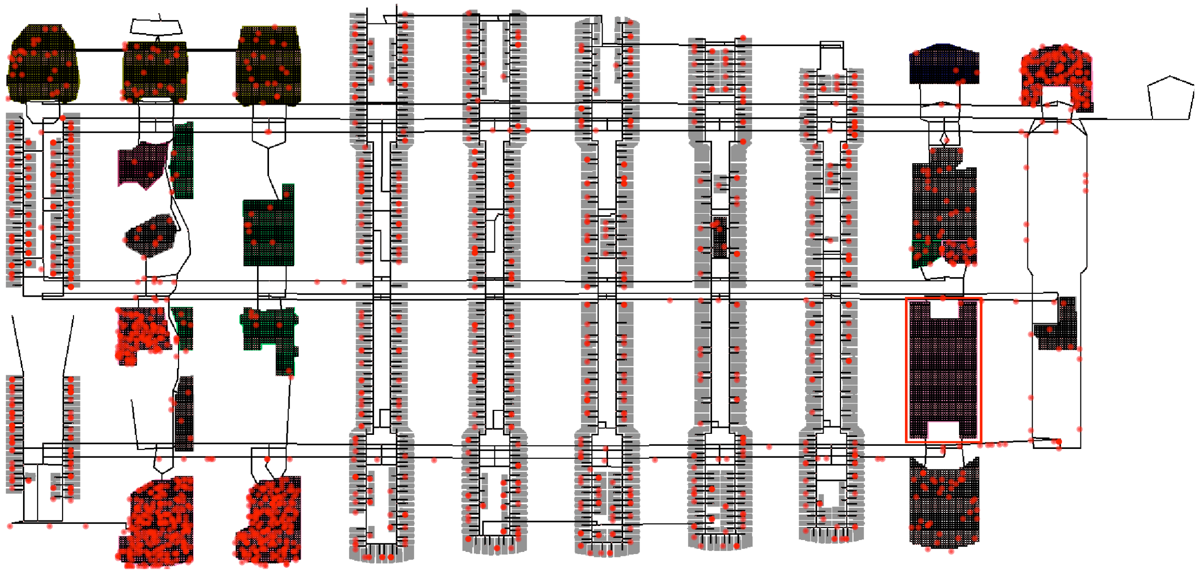

408 **Supplementary Figure S9.** Simulations for Norovirus outbreak with Dining Restricted protocol. The figure shows the rise in infections when one of the main dining areas is closed (shown here in red outline). This is due to overcrowding in the other dining rooms.

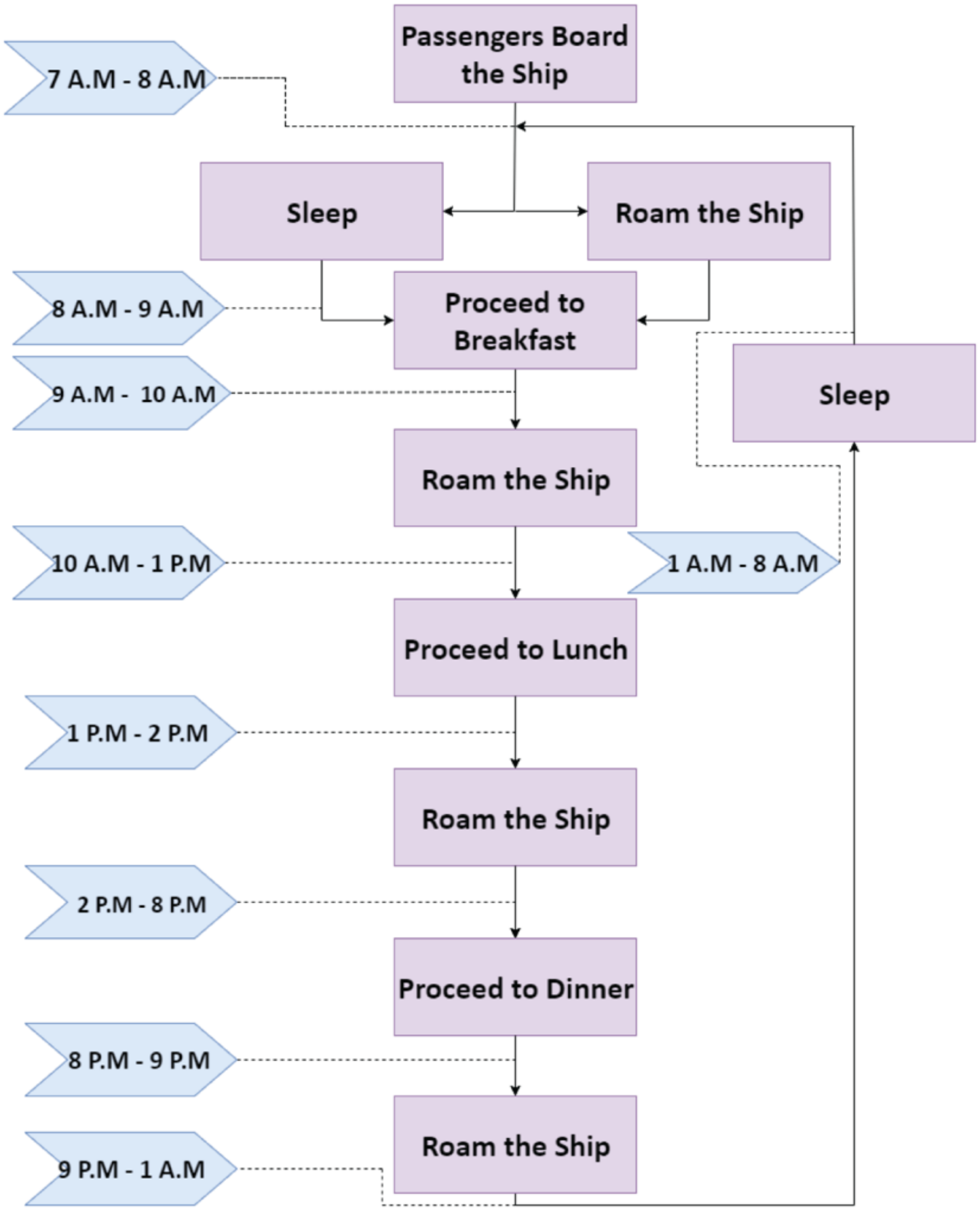


**Supplementary Figure S10.** Flowchart depicting a summary schedule for an intelligent agent modeled as a passenger. Stochasticity is introduced in the recreational activities that a passenger agent performs.

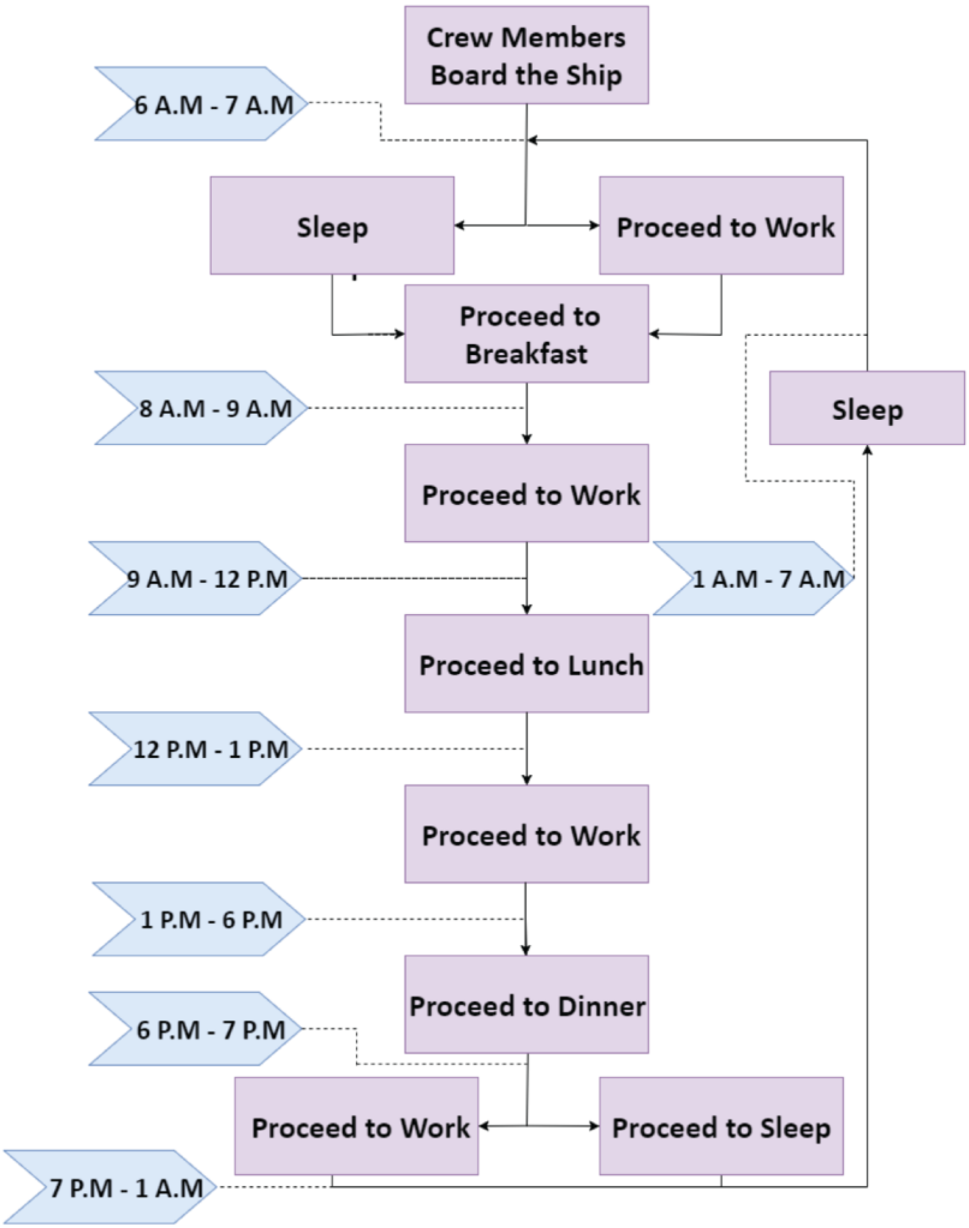


**Supplementary Figure S11.** Flowchart depicting a summary schedule for an intelligent agent modeled as a crew member. The crew agents spend most of their time in particular areas of the cruise ship performing tasks.

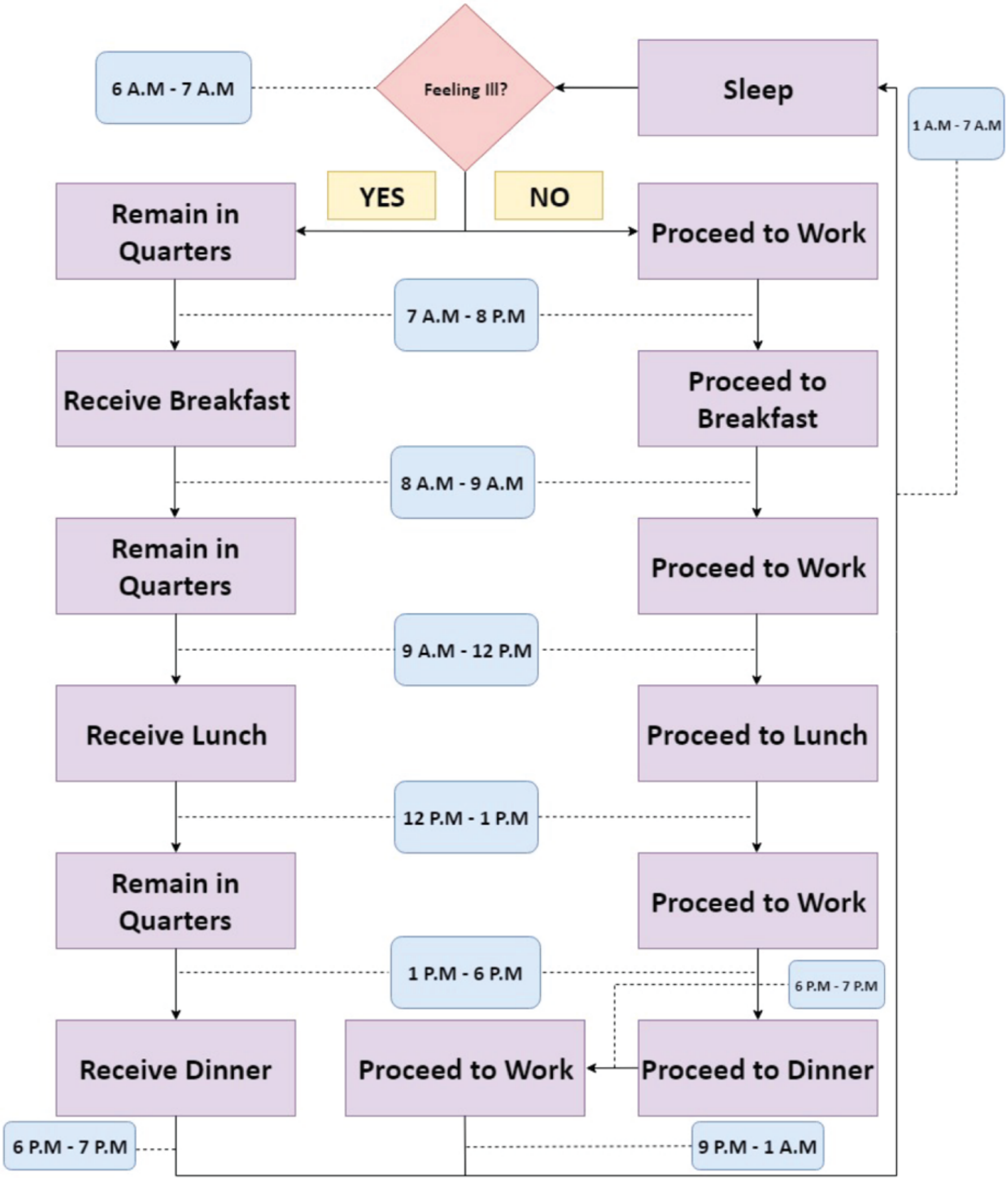


**Supplementary Figure S12.** Flowchart depicting a summary schedule for an infected and ill crew agent who is following containment measures. The crew member continues to remain in quarters until recovered.

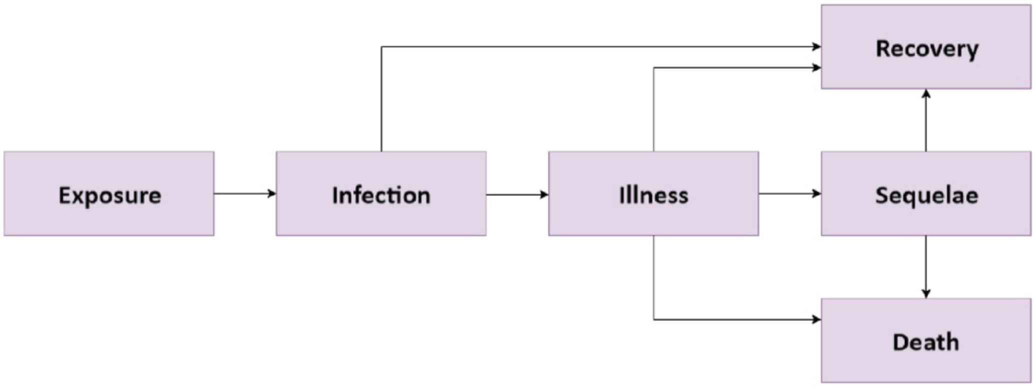


**Supplementary Figure S13 .** The major phases of any infectious disease that can be modeled for an agent. The pathogens with consequential fatality rate, such as SARS-CoV-2 and EBOV have modeling for the Death phase.

### References


1. S. Luke, C. Cioffi-Revilla, L. Panait, K. Sullivan, G. Balan, Mason: A multiagent simulation environment. *Simulation* **81**, 517-527 (2005).
2. M. Loy, R. Eckstein, D. Wood, J. Elliott, B. Cole, *Java swing*. (" O'Reilly Media, Inc.", 2002).
3. C. N. HAAS, Estimation of risk due to low doses of microorganisms: a comparison of alternative methodologies. *American journal of epidemiology* **118**, 573-582 (1983).
4. P. Teunis *et al.*, Shedding of norovirus in symptomatic and asymptomatic infections. *Epidemiology & Infection* **143**, 1710-1717 (2015).
5. D. K. Ip *et al.*, Viral shedding and transmission potential of asymptomatic and paucisymptomatic influenza virus infections in the community. *Clinical infectious diseases* **64**, 736-742 (2017).
6. T. P. Weber, N. I. Stilianakis, Inactivation of influenza A viruses in the environment and modes of transmission: a critical review. *Journal of infection* **57**, 361-373 (2008).
7. R. L. Atmar *et al.*, Norwalk virus shedding after experimental human infection. *Emerging infectious diseases* **14**, 1553 (2008).
8. P. Liu *et al.*, Persistence of human noroviruses on food preparation surfaces and human hands. *Food and Environmental Virology* **1**, 141-147 (2009).
9. P. Teunis, A. Havelaar, The Beta Poisson dose ‐ response model is not a single ‐ hit model. *Risk Analysis* **20**, 513-520 (2000).
10. P. F. Teunis *et al.*, Norwalk virus: how infectious is it? *Journal of medical virology* **80**, 1468-1476 (2008).
11. P. F. Teunis, N. Brienen, M. E. Kretzschmar, High infectivity and pathogenicity of influenza A virus via aerosol and droplet transmission. *Epidemics* **2**, 215-222 (2010).
12. R. A. Blair, B. S. Morse, L. L. Tsai, Public health and public trust: Survey evidence from the Ebola Virus Disease epidemic in Liberia. *Social science & medicine* **172**, 89-97 (2017).
13. C. L. I. Association. (2018).
14. V. J. Munster *et al.*, Pathogenesis and transmission of swine-origin 2009 A (H1N1) influenza virus in ferrets. *Science* **325**, 481-483 (2009).
15. G. P. Kobinger *et al.*, Replication, pathogenicity, shedding, and transmission of Zaire ebolavirus in pigs. *Journal of Infectious Diseases* **204**, 200-208 (2011).
16. K. K.-W. To *et al.*, Consistent detection of 2019 novel coronavirus in saliva. *Clinical Infectious Diseases* **71**, 841-843 (2020).
17. A. Sakurai *et al.*, Natural history of asymptomatic SARS-CoV-2 infection. *New England Journal of Medicine* **383**, 885-886 (2020).
18. M. M. Arons *et al.*, Presymptomatic SARS-CoV-2 infections and transmission in a skilled nursing facility. *New England journal of medicine* **382**, 2081-2090 (2020).
19. G. Onder, G. Rezza, S. Brusaferro, Case-fatality rate and characteristics of patients dying in relation to COVID-19 in Italy. *Jama* **323**, 1775-1776 (2020).
20. C. f. D. C. a. Prevention. (2024).
21. L. Huang *et al.*, Rapid asymptomatic transmission of COVID-19 during the incubation period demonstrating strong infectivity in a cluster of youngsters aged 16-23 years outside Wuhan and characteristics of young patients with COVID-19: A prospective contact-tracing study. *Journal of infection* **80**, e1-e13 (2020).
22. J. Papenburg *et al.*, Household transmission of the 2009 pandemic A/H1N1 influenza virus: elevated laboratory-confirmed secondary attack rates and evidence of asymptomatic infections. *Clinical Infectious Diseases* **51**, 1033-1041 (2010).

23. S. Judson, J. Prescott, V. Munster, Understanding ebola virus transmission. *Viruses* **7**, 511-521 (2015).
24. P. Mlcochova *et al.*, SARS-CoV-2 B. 1.617. 2 Delta variant replication, sensitivity to neutralising antibodies and vaccine breakthrough. (2021).
25. M. W. Fong *et al.*, Nonpharmaceutical measures for pandemic influenza in nonhealthcare settings—social distancing measures. *Emerging infectious diseases* **26**, 976 (2020).
26. C. f. D. C. a. Preventation. (2024).
27. E. H. Cramer, C. J. Blanton, C. Otto, V. S. P. E. H. I. Team, Shipshape: sanitation inspections on cruise ships, 1990–2005, vessel sanitation program, centers for disease control and prevention. *Journal of Environmental Health* **70**, 15-21 (2008).
28. U. R. Bodnar *et al.*, Preliminary guidelines for the prevention and control of influenza-like illness among passengers and crew members on cruise ships. (1999).
29. L. Fu *et al.*, Clinical characteristics of coronavirus disease 2019 (COVID-19) in China: a systematic review and meta-analysis. *Journal of Infection* **80**, 656-665 (2020).
30. L. Pan *et al.*, Clinical characteristics of COVID-19 patients with digestive symptoms in Hubei, China: a descriptive, cross-sectional, multicenter study. *Official journal of the American College of Gastroenterology| ACG* **115**, 766-773 (2020).
31. J. T. Brooks, J. C. Butler, Effectiveness of mask wearing to control community spread of SARS-CoV-2. *Jama* **325**, 998-999 (2021).
32. C. J. Willmott, K. Matsuura, Advantages of the mean absolute error (MAE) over the root mean square error (RMSE) in assessing average model performance. *Climate research* **30**, 79-82 (2005).
33. F. J. Massey Jr, The Kolmogorov-Smirnov test for goodness of fit. *Journal of the American statistical Association* **46**, 68-78 (1951).
34. J. Shaman, A. Karspeck, Forecasting seasonal outbreaks of influenza. *Proceedings of the National Academy of Sciences* **109**, 20425-20430 (2012).
35. W. O. Kermack, A. G. McKendrick, A contribution to the mathematical theory of epidemics. *Proceedings of the royal society of london. Series A, Containing papers of a mathematical and physical character* **115**, 700-721 (1927).
